# Mechanical load conditions the spectrin network to ‘runon’ proteolysis and promotes early onset neurodegeneration

**DOI:** 10.1101/2024.11.04.621798

**Authors:** Nawaphat Malaiwong, Anne-Kristin Dahse, Li-Chun Lin, Ravi Das, Montserrat Porta-de-la-Riva, Nicole Scholz, Michael Krieg

**Affiliations:** ICFO - Institut de Ciències Fotòniques, The Barcelona Institute of Science and Technology, Castelldefels (Barcelona) 08860, Spain; Rudolf Schönheimer Institute of Biochemistry, Division of Molecular Biochemistry - Cellular Neuromechanics and Adhesion GPCR group, Medical Faculty, Leipzig University, Johannisallee 30, 04103 Leipzig, Germany

## Abstract

The correct homeostasis of the neuronal cytoskeleton and its dynamics is important for health and disease. Forces constantly act on the neurons in our body, leading to subtle axonal deformations and length changes. The spectrin cytoskeleton is known as a key player that protects neurons against mechanical damage. How the spectrin cytoskeleton changes with age and how it influences mechanoprotection in aging animals is not well understood. Using an interdisciplinary approach, we show that age causes a loss of proprioception during the first few days of adulthood in *C. elegans* via spectrin unfolding, loss of mechanical tension and degradation of the spectrin cytoskeleton. Guided by a proteomic screen to identify potential spectrin binding partners, we found that this early-onset neurodegeneration can be suppressed in *clp-1* mutants and by targeted expression of an engineered chaperone derived from human *α*B-crystallin. Our data suggest that the spectrin cytoskeleton is sensitized to proteolytic damage by calcium-sensitive proteases when mechanical stresses conspires with high-calcium concentrations as in proprioceptive signaling. These results may have implications for the etiology of diseases in which high calcium dynamics and mechanical stress co-incide.

## 1 Introduction

The decline in neuronal function is a prominent characteristic of natural aging. However, the precise connection between the molecular, cellular, and ultrastructural changes, the modifications in neuronal dynamics and signaling, and the resulting impairments in behavior that arise in advanced age remain uncertain. Even when significant neuronal loss is not typically observed at the cellular level in the course of early aging in healthy humans^1^ and genetic models^2–4^, alterations in excitability and signaling have been documented^1^. Many of these deficits relate to changes in sensory functions and processing^5^, implying early onset neurodegeneration. One of the earliest visible symptoms of age in adults is a progressive decline in body coordination and proprioception^6^, which can massively affect overall motor function and thereby quality of life. Understanding the mechanistic underpinnings of neurodegenerative processes, and particularly how aging influences the mechanobiology of neurons, holds immense potential for novel therapeutic targets and interventions^7^. However, mechanistic insight underlying age-dependent loss of proprioceptive functionality is scarce. This is due in part to the convergence of several cell-autonomous and non-autonomous signaling routes that collectively shape age-related alterations in cell mechanics and mechanosensation. For example, recent work in *C. elegans* has shown that cuticle collagens and apical extracellular matrix (ECM) stiffness change during age and that mechanotransduction in the epidermis can slow aging through ECM remodeling^8^. These data highlight the importance of the age-related changes of the neuronal mechanical properties and those of the surrounding tissues for optimal functionality of mechanosensory processes and mechanotransduction^9^.

In mammalian neurons, age and the exposure to mechanical stress correlates with structural changes in axon morphology^10^ and accumulation of proteolytically cleaved proteins^11^. Such cleavage events occur in the spectrin cytoskeleton and are evident in the formation of protein aggregates. Strikingly, calpain (*clp*)-mediated proteolytic products of the cytoskeleton^12^, including the fragmented spectrin network, are established cerebrospinal fluid-based biomarkers for neurodegenerative diseases, secondary to severe traumatic brain injury^13, 14^, and age^15^. This provides a plausible link between the integrity of the cytoskeleton, mechanical stress and the decline of neuronal function during aging. However, why proteolysis differentially affects neurons in old animals and the factors leading up to the aggregation are not resolved.

The spectrin network is a ubiquitous component of the cytoskeleton that protects axons of moving animals against mechanical stresses^16, 17^. It consists of periodically located *α*/*β* heterotetramers that provide mechanical support to the plasma membrane through linkage with the actin and microtubule cytoskeleton^18^. Despite the ubiquitous appearance of the 190 nm periodic pattern of the spectrin network in cultured neurons^17, 19, 20^, the spectrin cytoskeleton has shown to be very dynamic *in vivo*^21, 22^, and can exist in an extended, but also compressed configuration^23^ depending on the mechanical stress level^24^.

Here, we use *C. elegans* as a model to demonstrate how mechanical stress and age collaborate as risk factors in early onset decline in neuronal function. Behavioral analysis and calcium imaging show that proprioceptive decline is one of the first ‘symptoms’ of aging in young adults. This deficit originates in a single neuron that fails to properly respond to changes in body posture. By leveraging a genetically encoded tension sensor, we demonstrate that the constitutive tension profile of the spectrin network within this neuron declines with age. Further, we identified the calcium-sensitive protease *clp-1* as a mediator in spectrin breakdown and demonstrate that loss of muscle contraction in paralysed animals or overexpression of human *α*B crystallin, a constitutive chaperone, suppresses spectrin cleavage and age-related defects in proprioception, respectively. Together, our data suggest a model in which high calcium signaling and mechanical unfolding of the spectrin cytoskeleton conspire in early-onset neurodegeneration, a phenomenon likely to be conserved across diverse biological systems.

## 2 Results

### Loss of proprioception as an early aging phenotype

To understand how age affects proprioceptive coordination during locomotion, we recorded the natural crawling behavior of individual *C. elegans* animals for 5 min on food and in the absence of external stimuli during the first 14 days of adulthood. With the aim of maintaining a synchronized population, cohorts of young adults (YA) were transferred to plates containing FUDR, which is a potent S-phase inhibitor that controls population size by inhibiting reproduction (^25^, Methods). We then used a previously established analysis pipeline for dimensionality reduction ^23, 26^ to display the behavior of the animals in their corresponding eigenworm space. Interestingly, animals that completed their second day of adulthood (2DA) already showed a significantly different behavior than young adults, visible as a larger diameter manifold in their 3D eigenworm space, indicative for deeper body bends during forward locomotion (Fig. 1a). We thus reasoned that these early defects resulted at least in part from a failure to curb muscle contraction, an indication for proprioceptive deficits ^23, 27^. This phenotype is also apparent after 4DA, but begins to disappear in later age, during which the behavioral fingerprint in the eigenworm space is insignificant compared to YA animals (Fig. 1b). Because animals generally moved less during these midlife ages, we reasoned that the transition involved general neurodegenerative and muscular defects. Indeed, after ten days, adult animals experience a rapid decline in locomotory activity associated with a catastrophic change in their behavioral fingerprint (Fig. 1b). Even though animals retain the ability to move after 6DA, previous results show that, alterations in the neuromuscular junctions and sarcopenia, govern the dramatic decline in the locomotion behavior in older animals^28, 29^. Importantly these changes were not affected by the presence of FUDR (Fig. S1a-d). Because the increase in body curvature did not depend on the canonical daf-2 insulin longevity pathway^30^ (Figure S1e,f), we reasoned it is independent on genetic factors that commonly lead to lifespan extension and reduction, but may be due to epigenetic and physiological effects. Aging related damage products accumulate stochastically and thus contributes to considerable variability of phenotypes ^31^. Short-lived animals have an earlier decline in lifespan compared to long-lived animals, which contributes to a large variability when comparing a trait at a chronological age. To estimate if the increase in body curvature is indeed caused by age-related effects, we re-scaled age to the lifespan of the animals. This analysis showed that the increase in the eigenworm space scales robustly according to their relative age (Fig. S2). Taken together,*C. elegans* exhibits locomotion defects early in aging, characterized by significant increases in body bending amplitude.

**Fig. 1.**
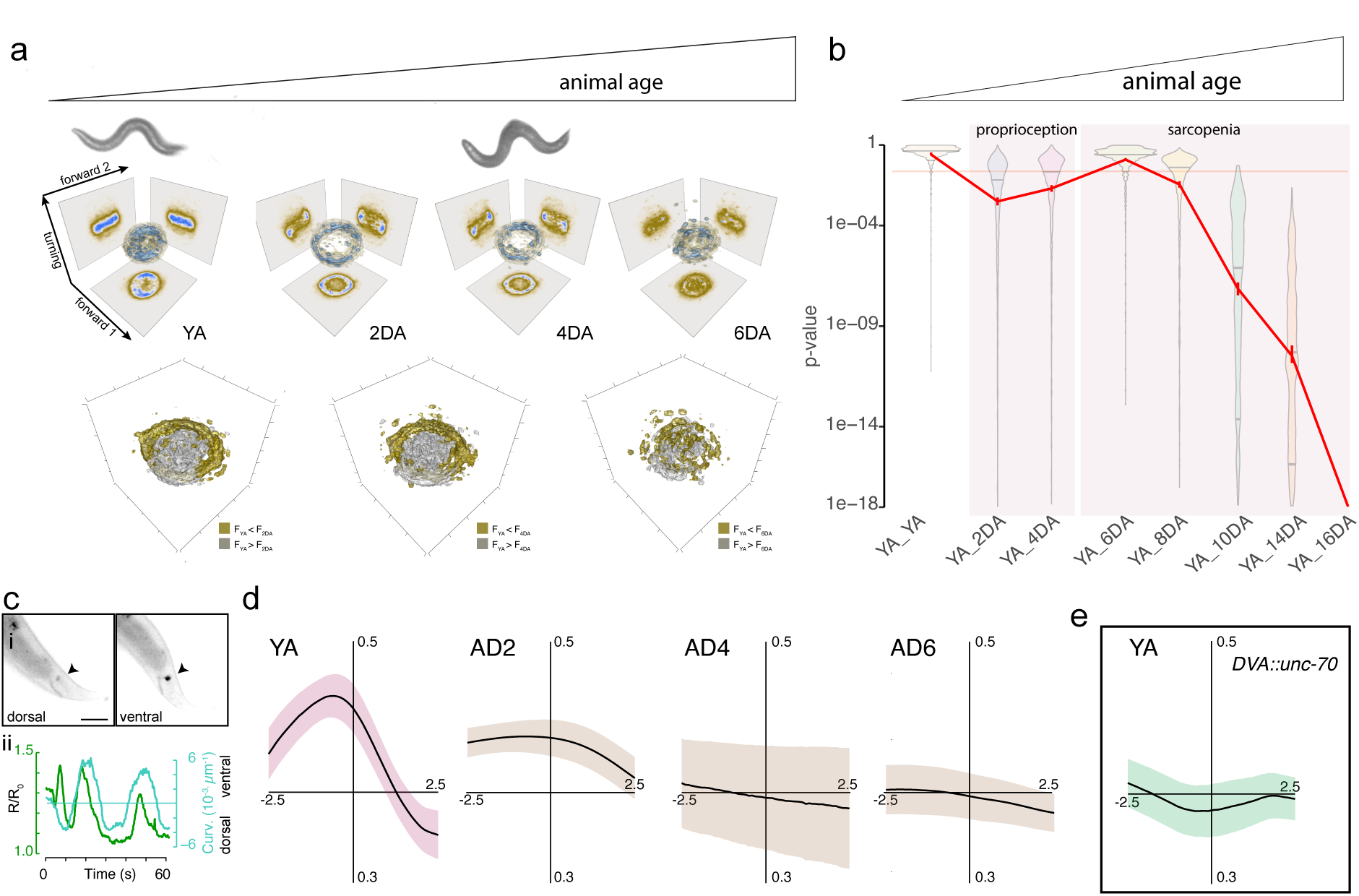
Functional decline in proprioceptive sensitivity leads to early age locomotion defects. **a,** Animal locomotion changes during early days of adulthood. Representative animal micrograph on top of the 3D density estimate for the joint probability distribution (equivalent to a discrete 3D histogram) of the two forward and turning modes in the eigenworm space. Color scale = (brown, low density; blue, high density). 3D voxelgram shows statistically significant differences in the probability densities between young adults and animals at an age that is indicated in the figure. Voxels in silver indicate larger density of young adults, voxels in gold indicate smaller than the aged animals. The test is performed on each voxel in a 100x100x100 grid (see Methods) and color coded according to its outcome using a two-sided Wald test. Voxels with p*>*0.01 are not shown. **b,** Violin plots of the *p*-value distributions for 1000 independent tests of a bootstrapped population estimate of wt 3D probability density function (ctrl) tested against itself or a bootstrapped population estimate of the aged sample. Orange line indicates *α*=0.05 level of significance for the hypothesis H_0_ that bootstrapped density functions derived from N2 and age animals are equal (see Methods). The red line indicates the mean p-value. The reproductive period ends when sarcopenia and muscle weakness sets in (4 day adults). **c,** i) Representative images of a young adult animal with GCaMP expression in DVA under different body postures. ii) Calcium intensity (green) and body curvature (turquoise) vs time. Scale bar = 40 µm. See also Suppl. Video 1. **d, e,** Cross correlation analysis of the calcium activity vs curvature in DVA for (d) wild type, young adults and progressively aged animals until day 6 and (e) *DVA:unc-70* mutant young adult animals.

### DVA calcium signals change during early aging

An increase in body curvature during forward crawling is a typical sign of proprioceptive impairment, and various neurons contribute to it^32^. We have previously shown that the proprioceptive interneuron DVA functions to limit body posture amplitudes in a spectrin-dependent manner^23^. In that process, DVA senses changes in body posture (through changes in axonal tension and compression) and responds with increased calcium activity during body bends. To better understand whether DVA activity is affected in older animals, we recorded calcium transients using a DVA-specific GCaMP indicator in partially restraint worms during dorso-ventral body bends. We found that DVA preferentially activated during ventral body bends (Fig. 1c i), consistent with the previous finding that DVA acts as a compression sensor^23^. During aging, the correlation between DVA activity and ventral body bends declined rapidly until 4DA, during which we occasionally observed ventral DVA calcium activity (Fig. 1c ii). The majority of the recordings, however, were uncorrelated to a specific body posture (Fig. 1d). We have obtained a similar qualitative situation in YAs that carried a DVA neuron-specific defect in *unc-70 β*-spectrin (Fig. 1e). Based on the similarity of the age-related symptons and the *β*-spectrin loss of function phenotype, we speculated that loss in DVA calcium activity and, concomitant defects in locomotion behavior stem from *β*-spectrin-related malfunction.

We and others have previously shown that a loss of *β*-spectrin function leads to severe buckling defects in TRNs and DVA during ventral bending^17, 22, 23, 33, 34^. We reasoned that a similar phenotype may appear in old animals. Testing this hypothesis, we imaged the morphology of DVA axons in YA and AD8 animals in different body postures. As expected YA axons did not display any major deformations during ventral bends^23^. The axons of old animals, however, appeared morphologically altered when ventrally, but not dorsally bend (Fig. 2a). This phenotype is strikingly similar to that of DVA^23^ or TRNs^22^ in *unc-70*/ *β*-spectrin loss-of-function animals. Collectively, these findings imply that defects in *β*-spectrin may contribute to a loss of mechanical stability in old animals.

**Fig. 2.**
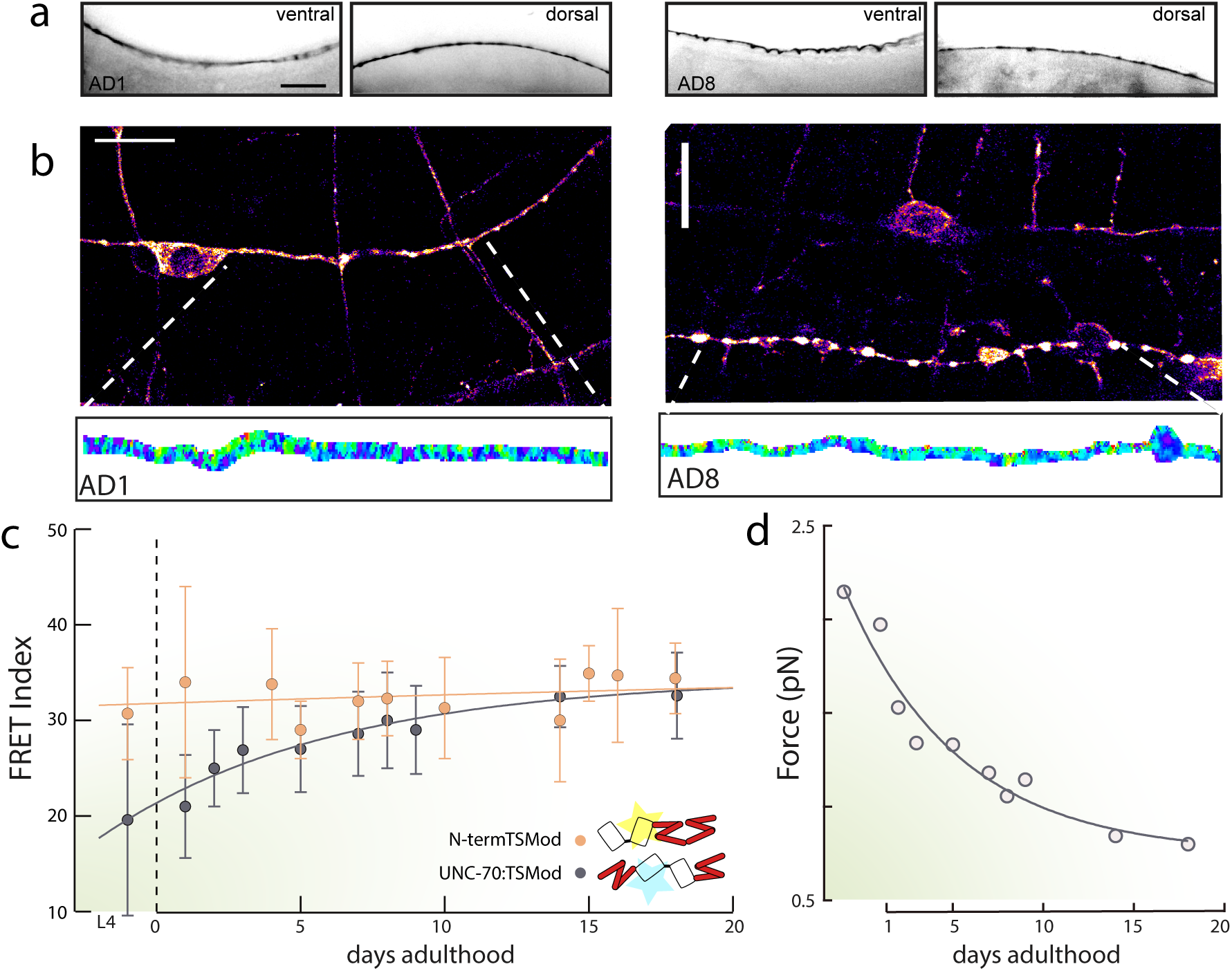
Tension in the spectrin network successively declines with animal age. **a,** Representative image of a DVA axon close to the cell body in a (i) young adult and an (ii) eight day old (8DA) animal during ventral and dorsal bends. Scale bar = 10 µm. **b,** Representative FRET images and the FRET map for young adults and a eight day adult animal. Scale bar = 5 µm. **c,** FRET index vs animal age of the unc-70:TSMod tension sensor and the force-insensitive control, mean± SD. Solid lines indicate fit to a single exponential function. **d,** Plot of the converted force vs animal age. Solid line is a best fit to a stretched exponential function (*f*(*t*) = *a* · exp(*τ t^b^*)).

### Spectrin tension declines with loss of proprioception

To further substantiate *β*-spectrins role in the age-related proprioceptive phenotype, we imaged the subcellular distribution and mechanical properties of the spectrin cytoskeleton utilizing a *β*-spectrin-based mechanical tension sensor (Fig. 2b and ref ^22, 35^). Consistent with earlier reports, we found that *β*-spectrin is under constitutive mechanical tension in L4 larval stages and AD1 animals (Fig. 2c, d). Surprisingly, we observed an increase in FRET efficiency (loss of tension) already in two day old adults, which progessively converged to the FRET values derived from a no force control (N-terminally tagged *β*-spectrins) at AD10 (Fig. 2b). Using the published conversion of the FRET to force values^36^, we calculated that the average force on *β*-spectrin dropped from 2.2 pN to less than 0.7 pN at 17DA (Fig. 2d). This indicates that *β*-spectrin undergoes a gradual decrease in mechanical tension during the aging process. Interestingly, this loss coincides with the beginning of visible *β*-spectrin aggregates along the neurites (Fig. S3a). In older animals, these aggregates became larger but less abundant, reminiscent of the growths of liquid-like condensates (Fig. S3b). To corroborate that these aggregates were not the result of an artefact due to the multi-copy transgene, we imaged animals expressing UNC-70::GFP and SPC-1::GFP tagged at their respective endogenous locus (Fig. S3c, d; ^37^). We observed a subtle but consistent increase in aggregated spectrin in distinct foci of various unidentified neurons in aging animals in both transgenic situations. This finding is in line with the recent suggestion that topological transitions in *β*-spectrin lead to biomolecular condensation and loss of spectrin tension in tissue culture cells ^38, 39^. Taken together, this shows that age directly affects the organization and tension values of the spectrin cytoskeleton, thereby controlling behavioral activity and body posture.

### Spectrin unfolding buffers mechanical stress *in vivo*

The stereotypic 190 nm periodicity of the spectrin cytoskeleton seen in super-resolution images of neurons in culture^17, 19, 20, 40^ suggests that the spectrin network exists in a fully extended state close to the natural contour length expected for an *α/β* spectrin heterodimer ^41^. However, in intact animals the spectrin periodicity seemed to be more variable (Fig. 3a, b), visible as multiple peaks in the Fourier analysis corresponding to 160 nm and 190 nm of the observed *β*-spectrin pattern in motorneurons (Fig. 3c i). We hypothesized that periodicity of *β*-spectrin is due to its flexibility and hence coupled to its intrinsic mechanical state. This notion is in agreement with reports demonstrating a reduction of periodicity in neurons with lower mechanical tension^17^. To understand whether the hypothetical difference in *β*-spectrin topology in differently oriented axons is due to differences in mechanical tension, we built a *β*-spectrin-based FRET-force sensor exclusively expressed in the retrovesicular ganglion and some motorneurons, including DA9 (*mig-13p*, see Methods and ref. ^42^). We then measured FRET in circumferential commissures and longitudinal axons independent of variability in cellular identity. Intriguingly, we found that FRET calculated from images of commissures was higher than FRET calculated on parts of the longitudinal neuron (Fig. 3c ii). This suggests that neuronal orientation and position within the animal may influence their mechanical state.

**Fig. 3.**
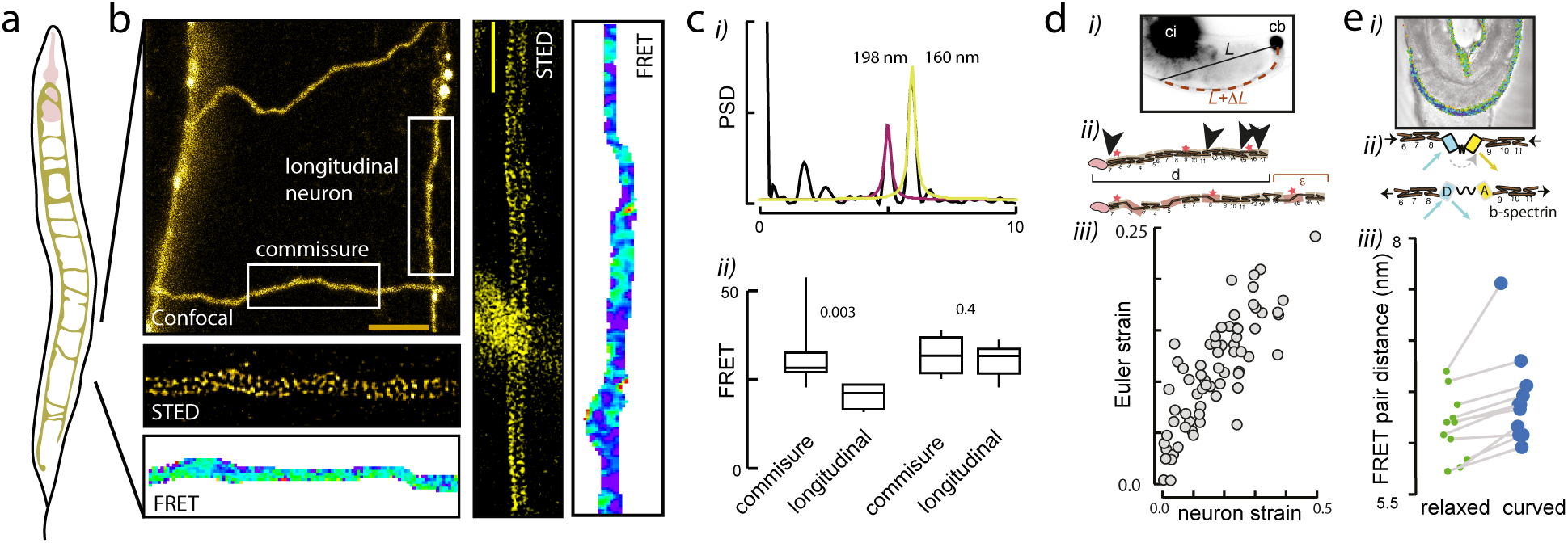
Spectrin buffers tension by unfolding individual repeats. **a,** Schematic of an animal with their commissures and longitudinal neurons. **b,** Confocal image showing commissures and longitudinal neurons. Scale bar = 5µm. Insets show representative superresolved ROI acquired with STED microscopy and a FRET index map derived from the UNC-70::TSMod. Scalebar = 2µm. **c,** Quantification of i) a power spectral density (PSD) of the intensity profile of the commissures shown in b. Multiple peaks appear corresponding to 160 and ≈ 190 nm. ii)FRET index for the UNC-70::TSMod tension sensor and force insensitive control acquired on commissures and longitudinally aligned neurites. p-value derived from a t-test. **d,** Spectrin unfolding accommodates neurite length changes during body bending. i) Snapshot of a DVA axon under dorsal body bend, leading to an extension of the neurite. L=resting length of the neuron in a straight animal; *ɛ*=normalized neuron length change (Δ*L/L*strain). ii) Schematic of the resting contour length and the extended spectrin length after unfolding. Arrowheads (black) point to predicted proteolytic cleavage sites; yellow arrowhead points to one of many cleavage sites identified in recombinant *β*II-spectrin^84^; red stars depict predicted unfolded domains. d=resting length of the spectrin tetramer in straight neurons; *ɛ*= change in length due to unfolding. iii) Plot of the calculated bending (Euler) strain (curvature x thickness) vs the measured neurite extension (Δ*L*). **e,** Unfolding buffers tension in the network. i) Snapshot of the FRET map overlayed with a brightfield image of an animal inside the waveform sampler microfluidic chip at high body bend. Image reproduced from Ref.^23^. ii) Schematics of the UNC-70::TSMod under extension. iii) Plot of the chromophore separation, calculated from the bleed through corrected FRET efficiency in the relaxed and bent state of the animal for N=15 animals.

Next, we wondered how animal movement might influence *β*-spectrin mechanics and flexibility. Because we were not able to quantify *β*-spectrin periodicity using STED superresolution in live, moving animals, we resorted to an indirect measurement. We have previously shown that TRN and DVA neurites elongate and shorten during locomotion, in response to dorsal or ventral body bends, respectively ^17, 23^. This strain Δ*L/L* is proportional to the distance of the neuron from the body center line^17^ and the curvature of the bend (Fig. 3d i) and can be more than Δ*L/L* = 40% of the resting length of the neuron (e.g. when the animal is not curved). An obvious question is, where does the additional material come from to accommodate the length change of several tens of a micrometer? Even though this increase in local length could be explained by incorporation of free *β*-spectrin molecules into the network, this assembly/disassembly of the *β*-spectrin cytoskeleton seems unlikely to occur within the time required to change body bends (*<* 2s). Our previous observation suggesting that UNC-70 experiences elevated mechanical tension and compression during body bends ^23^, together with spectrin’s proposed conformational flexibility ^43, 44^ pointed at the possibility that local unfolding of individual spectrin domains enabled the curvature-associated increase in neuronal length during animal locomotion (Fig. 3d ii). In fact, *β*-spectrin was shown to unfold under physiological mechanical stress in red blood cells^21^ and isolated chick neurons^24^ when experiencing mechanical strain of *<*30%, a value similar to strain values that occur during normal locomotion in *C. elegans*^17^. To better understand if axonal *β*-spectrin indeed unfolds during normal locomotion, we measured the curvature-dependent axon elongation and mechanical tension in *β*-spectrin from data obtained in ^17^ and ^23^ to specifically evaluate the tension-strain relationship (Fig. 3d iii). The maximum decrease in FRET efficiency of 15% at the highest curvatures in the microfluidic chip suggests an elongation of *<*8 nm (Fig. 3e iii). Given the overall length of the spectrin dimer of 90 nm, this would indicate a strain of *ε <*10%. This is 4 times lower than the extension Δ*L* of the axons, because we measure axon strains at the same curvature of about 40% in wild type animals. We therefore conclude that tension within the spectrin network is buffered and the additional elongation occurs due to transient unfolding of individual spectrin repeats^21, 45–47^ such that 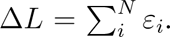 Taken together, our data support a model in which spectrin is highly dynamic *in vivo* and acts as a force buffer that constantly undergoes conformational changes through unfolding/refolding cycles.

### Proteolytic degradation of the spectrin network starts in early adulthood

Thus far we have shown that regular proprioception begins to change within the first days of adulthood accompanied by aberrant calcium activity in DVA as well as a drop in spectrin network tension and *β*-spectrin aggregation in neurons. In an attempt to understand how age relates to the loss of mechanical tension within the spectrin network, we started searching for putative interactors. To this end, we conducted UNC-70::GFP pull-downs from protein homogenisates of a synchronized population of wild type animals. Subsequent mass spectrometry, yielded many potential binding partners which included stereotypic interactions that involve the actin and microtubule cytoskeleton, but also components of the cellular proteostasis machinery such as chaperones, proteases and ubiquitin ligases (Supplementary Table 1, Fig. 4a) as candidates for potential *β*-spectrin interaction partners.

**Fig. 4.**
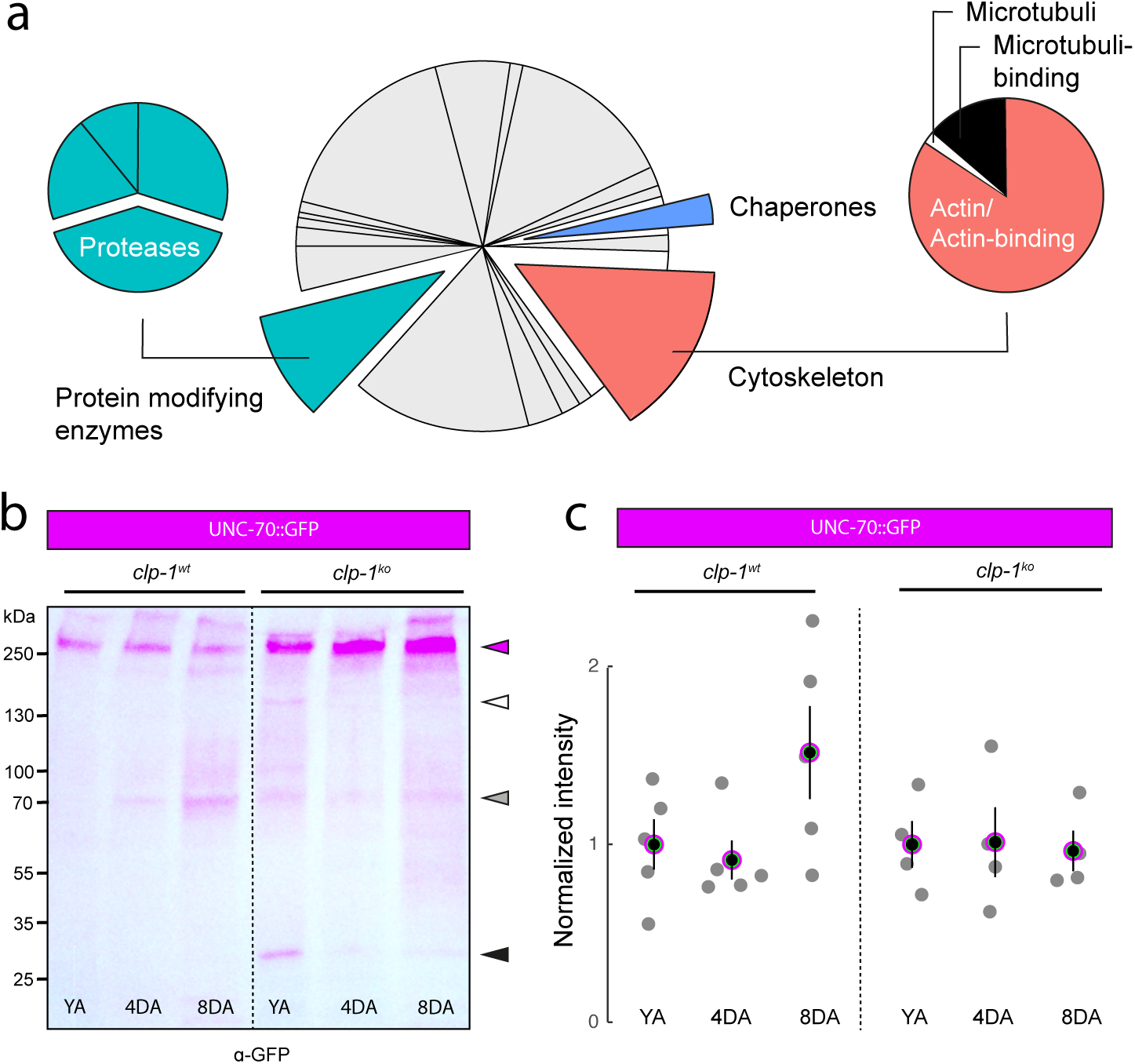
Spectrin is degraded by calpain-1 in aging animals. **a,** Categorization of the most relevant interaction partners obtained by mass spectrometry analysis. A comprehensive list can be found in Supplementary Table 1. **b,** Western blot against UNC-70::GFP in wild type and *clp-1* KO animals for age-matched samples. **c,** Quantification of the western blots. Red dot shows mean± standard error normalized to unstained N2 animals and the fold change to young adults (YA). Analysis was restricted up to 8DA due to the difficulty in getting enough material in agematched samples.

Among these candidates, the calcium-dependent cysteine-type endopeptidase calpain (CLP-1) was a candidate to consider. CLP-1 has previously been shown to cleave *β*-spectrin^48, 49^ in vertebrate brain tissue after traumatic injury^50^ and during age^51, 52^. Based on this published data and predictive models^48, 53–55^, we hypothesized that calpain cleaves and thereby degrades UNC-70.

To directly test whether spectrin gets degraded in a CLP-1 dependent manner, we performed western blot analyses of whole animal lysates derived from an age-synchronized population of YA, 4DA and 8DA animals (Fig. 4b, c, and Supplementary Fig. 4). In wild type YA, we repeatedly detected a 250 kDa band, which corresponds to the predicted molecular weight of GFP-tagged full length (FL) *β*-spectrin. When we performed the same analysis with protein lysates from older animals, where smaller fragments appeared in expense of the spectrin fulllength band (Fig. 4b, c). The largely undefined smear suggest that spectrin dies not get deggraded at a single site, but can have various, perhaps randomly accessible cleavage sites. To test whether this fragmentation of FL spectrin involves proteases and specifically CLP-1, we repeated the western blot analysis with protein samples from animals that were mutant for CLP-1 (*clp-1(mir60)*). Indeed, here we did not detect a decrease in the quantity of FL spectrin and smaller fragments were much less abundant (Fig. 4b, c). These observations strongly suggest an age- and CLP-1-dependent degradation of spectrin.

Next, we wondered if we can detect whether CLP-1 and UNC-70 directly interact with each other using co-immunoprecipitation assays. Using CRISPR, we generated a stable knock-in of a fluorescent protein at the endogenous *clp-1* (CLP-1:mKate) and pulled down UNC-70::GFP from protein lysates prepared from young adults animals using GFP-antibody. Even though we could observe both fusion proteins in the lysates before CoIP, i.e. in the supernatant, we were unable to visualize a direct CLP-1/UNC-70 interaction (Fig. S4). We thus conclude that CLP-1::mKate does bind very transiently to UNC-70::GFP, or requires additional factors such as the presence of mechanical tension which is absent in the pulldown assay from dissociated cells. Importantly, similar observations have been made with collagen and MMP-1^56^, focal adhesion kinase and calpain^57^ and van Willebrand factor and ADAMTS13^58^.

Together, the data suggest that CLP-1 is involved in age-dependent, mechanoenzymatic degradation of the spectrin network *in vivo*.

### Calpain activity in DVA augments age-related locomotion defects

Spectrin has several proposed calpain cleavage sites (Fig. 5a; ^53, 54, 59^) and is known as a steretypical target for calpain in mammals^48, 52, 60^. Based on our protein biochemical data on CLP-1, we were wondering about its effect on animal locomotion. We generated a mutant allele with abolished *clp-1* function and found a mild defect in the locomotion of YA (Fig. 5b) and a severe reduction in lifespan (Fig. S5a). *clp-1* has its peak expression in mechanosensory neurons such as PLM and DVA^61^, but it is also expressed in many, if not all, other cells (Fig. 5c). Hence, while it cannot be ruled out that the drastic decline in lifespan could have a mechanical homeostasis component, it is most likely the result of the faulty proteolysis/degradation machinery affecting a plethora of biological processes^62^.

**Fig. 5.**
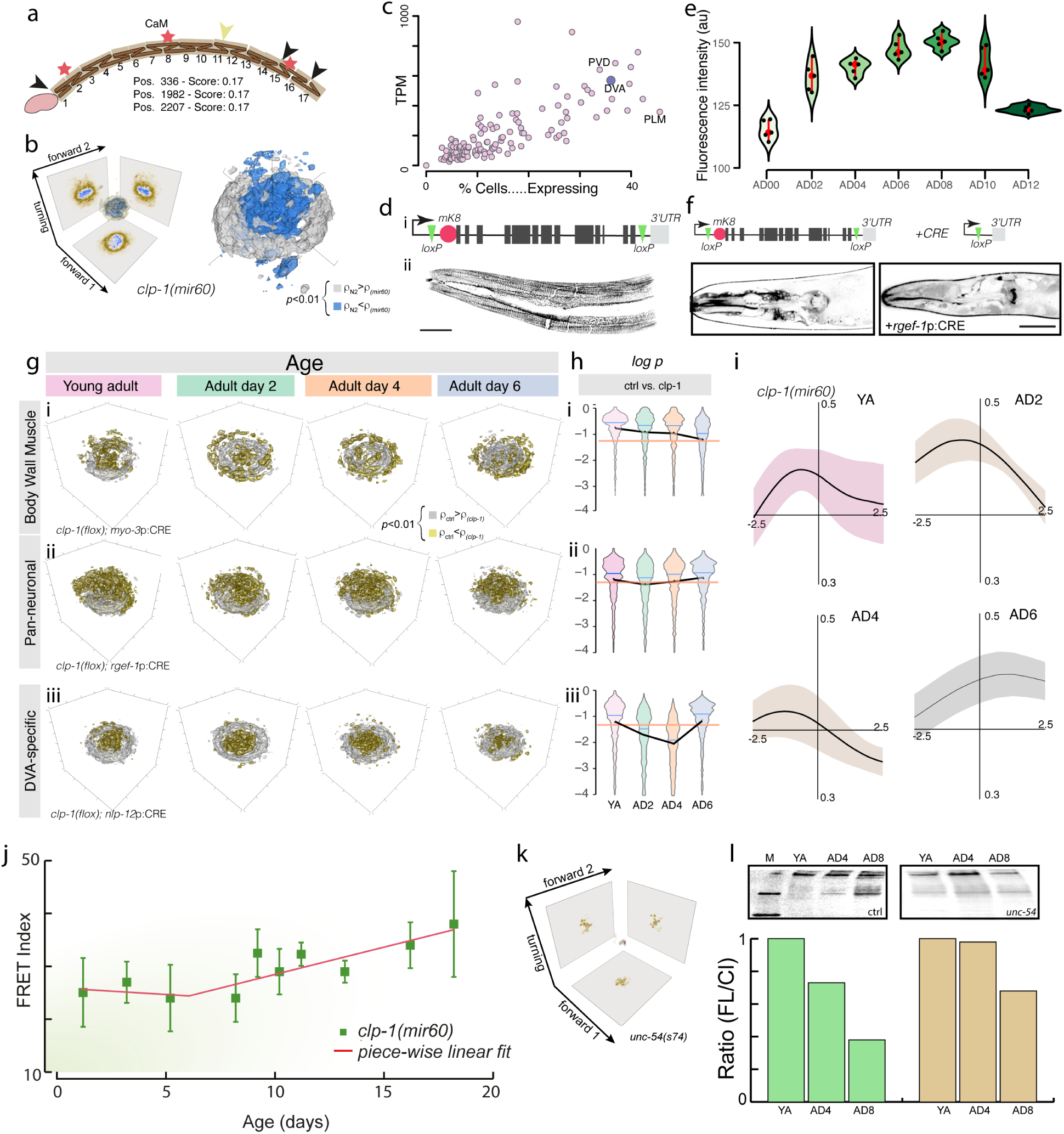
Calpain expression in DVA leads to proprioceptive decline. **a,** Secondary structure of *unc-70*/*β*-spectrin with the actin binding domain at the n-terminus and the 17 spectrin repeats. Red stars indicate sites with demonstrated force-induced conformational flexibility, black arrowheads indicate predicted Calpain cleavage sites. **b,** 3D plot of the estimate for the joint probability distribution (equivalent to a discrete 3D histogram) of the two forward and turning modes in the eigenworm space for the *clp-1(mir60)* mutation. Color scale = (brown, low density; blue, high density). Statistical test that the local kernel density function *ρ* of young adults is larger (blue) or smaller (grey) than the aged animals as shown in above. The test is performed on each voxel in a 100x100x100 grid (see Methods) and color coded according to its outcome. Voxels with p*>*0.01 are not shown. **c,** Quantification of *clp-1* transcripts in all neurons from ref. ^61^. **d,** i) Genomic organization of the *clp-1* locus with their lox sites and the location of the mKate tag. ii) Representative maximum intensity projection of a z-stack taken on the mKate::CLP transgenic animal, resembling the localization to the fibrous organelles (FO) in the epidermis. **e,** Quantification of the CLP-1:mKate intensity during age. Each black dot represents an animal; red point and vertical bar indicates median±95% confidence interval. **f,** Representative confocal images of the loxP-flanked *clp-1:mKate* transgenic animal and the loss of fluorescent signal in neurons after CRE expression under the *rgef-1* promoter (Scale bar = 40 µm). **g, h** Pan-neuronal and DVA-specific knockout of *clp-1* suppress the locomotion defect during early age. (g) 3D plot of the statistically significant differences comparing the joint probability distribution (equivalent to a discrete 3D histogram) of the two forward and turning modes in the eigenworm space for the tissue-specific *clp-1* mutations between wild type N2 and age-matched *clp-1* mutant with defects in i) all body wall muscles, ii) all neurons and iii) specifically in DVA at four different ages. Silver voxel indicates higher density for N2 control animal, golden voxels indicate higher density for *clp-1* mutants with p*<*0.01. (h) Violin plots of the *p*-value distributions for 1000 independent tests of a bootstrapped population estimate of control 3D probability density function of a N2 control animal tested against a bootstrapped population estimate of the age matched sample from the *clp-1* mutant. i) all body wall muscles, ii) all neurons and iii) specifically in DVA. Orange line indicates *α*=0.05 level of significance for the hypothesis H_0_ that bootstrapped density functions derived from N2 and aged animals are equal (see Methods). The blue line shows the mean p-value. The black line connects the median p=value. **i,** Cross correlation analysis of the calcium activity vs curvature in DVA for *clp-1(mir60)* young adults and progressively aged animals until day 6. **j,** FRET index vs animal age of the UNC-70:TSMod tension sensor in the *clp-1* mutant background, mean± SD. Solid lines indicate fit to a piece-wise linear function. **k,** 3D plot of the estimate for the joint probability distribution (equivalent to a discrete 3D histogram) of the two forward and turning modes in the eigenworm space for the *unc-54* mutation. **l** Representative western blot and their quantification of the UNC-70 full length/cleavage product ratio normalized to the young adult levels in the control and paralyzed *unc-54* mutation.

To further investigate the contribution of CLP-1 and the proteolytic cleavage of *β*-spectrin in the aging process, we used transgenic animals tagged with mKate at the endogenous *clp-1* locus and quantified their expression levels at different ages (Fig. 5d i). We identified expression predominantly in epidermis, muscles and neurons, including DVA (Fig. 5d ii). The expression gradually increased until 8DA followed by a sudden drop after 10DA (Fig. 5e). This indicates differential regulation of *clp-1* expression in different adult stages. Importantly, this data is supported by recent single cell RNA sequencing experiments showing that *clp-1* expression is increased in DVA until 5DA and begins to decline after 8DA (Fig. S5b, ^63^).

We next aimed to investigate tissue-specific effects of *clp-1* in locomotion and its involvement in aging. To do so, we equipped the strain with the endogenous mKate tag with two loxP sites and used previously characterized CRE drivers to delete clp-1 in a tissue specific manner^23^. We verified that the coexpression of a neuron-specific CRE leads detectable loss in mKate expression in neurons (Fig. 5f). We first examined the effect of *clp-1* knockout after expression of the CRE recombinase driven by a muscle-specific promotor and recorded 5 min videos every other day during the first week of adulthood. We found that animals lacking *clp-1* expression in the muscles crawled similar to age-matched wild type controls (Fig. 5g,h), which is also visible in the homogenous distribution of the density difference (mixed golden and silver voxels) in the 3D Eigenworm space (Fig. 5g).

To examine whether the loss of proprioception during early aging is due to loss of *clp-1* in neurons, we performed behavioral experiments using a pan-neuronal promoter. We found that animals crawled with a generally lower curvature (Fig. 5g ii and Supplementary Fig. 5c), but the age-related defect in proprioception is partially suppressed (Fig. 5h ii and Supplementary Fig. 5d), suggesting that *clp-1* has minor roles in the nervous system. To directly ask if *clp-1* has a neuron-specific role in DVA on locomotion, we deleted *clp-1* from DVA after expressing CRE under the cell-specific *nlp-12* promotor^23, 64^. Similar to the pan-neuronal knockout, though more robustly, we observed that the age-related proprioceptive decline was partially suppressed (Fig. 5g,h iii and Supplementary Fig. 5d). These data suggest that CLP-1 activity in DVA influences proprioceptive control of locomotion. To corroborate this conclusion, we next asked if calcium activity is altered in DVA of *clp-1* mutants and visualized calcium activity during dorso-ventral body bends (Fig. 5i). While wildtype animals show a pronounced de-correlation of calcium activity and body posture with age (Fig. 1), the calcium activity of *clp-1* mutant animals remained well correlated to the change in body posture in old animals.

This observation and the *in vitro* data on spectrin proteolysis bears the question if the age-related drop in spectrin tension can be delayed when CLP-1 is absent. We thus imaged UNC-70:TSMod in *clp-1* knockout background (up to 18DA) and found that the FRET values are generally more variable between individuals but also throughout life. While in wild type controls FRET values increased progressively starting after day two, i.e. spectrin tension relaxed (Fig. 2c), *clp-1* mutants retained constant FRET values until day seven, even though we measured higher FRET index values in general (Fig. 5j). After day eight, FRET values began to rise monotonically until the end of the experiment (18DA), suggesting a mechanism independent of CLP-1 that leads to loss of tension in the spectrin network. These data show that loss of *clp-1* partially rescues the age-dependent loss of spectrin tension.

Lastly, to understand if mechanical stress modulates UNC-70 degradation, we performed a western blot of UNC-70::GFP in the *unc-54* (encodes myosin heavy chain) background^65^. These animals barely move due to reduced muscle contraction and thereby exert less mechanical stresses on neurons and other body parts. A similar background was previously used to study the effect of self-inflicted forces on neuronal integrity^16, 66^. As indicated by the isolated densities (fix points) and the absence of a circular pattern in the eigenworm space, these animals were severely paralyzed (Fig. 5k). As expecetd, western blot analysis uncovered that loss of *unc-54* partially suppressed the degradation of UNC-70::GFP, visible as a reduction in the cleavage pattern (full length (FL) to cleavage products (CL)) compared to control animals (Fig. 5l). This observations point at the degradation of UNC-70 as a result of the continuous stretching to which it is exposed upon worm crawling.

Together, our data suggest that CLP-1 mediated proteolysis of *β*-spectrin in DVA promotes defective locomotion and calcium dynamics, leading to a loss of proprioceptive control and early onset neurodegeneration.

### Chaperones protect *β*-spectrin against mechanical damage

Exposed, hydrophobic sites are often recognized and protected by small heat-shock proteins of the *α*B-crystalin family^57^, which compete against enzymes that target these exposed cryptic binding sites for proteolysis^58^. Thus, engineered mini-chaperones have been proposed as therapeutic agents against conditional protein misfolding disorders^67^. Interestingly, HSP-25, an *α*B-crystallin homolog in *C. elegans*, has previously been shown to interact with *α*-actinin and vinculin, two components of dense bodies in muscles ^68^ (Fig. 6a). These dense bodies are similar to focal adhesions^69^, are subjected to myosin-dependent contractile forces and bear components that unfold in response to mechanical stress ^70, 71^ (Fig. 6a). It is thus plausible that chaperones also protect cytoskeletal elements of *C. elegans* neurons against aberrant proteolysis at transiently unfolded sites. Indeed, our mass-spectronomic analysis also identified HSP-25/*α*B-Crystallin (Supplementary Table 1) as a putative interactor. Intriguingly, as *hsp-25* expression is known to decline with age (Fig. S6a and ref.^63^), we hypothesize a loss in proteostasis which may give rise to uncurbed, ‘run-on’ degradation at mechanically sensitive moleculas through CLP-1 and other proteases.

**Fig. 6.**
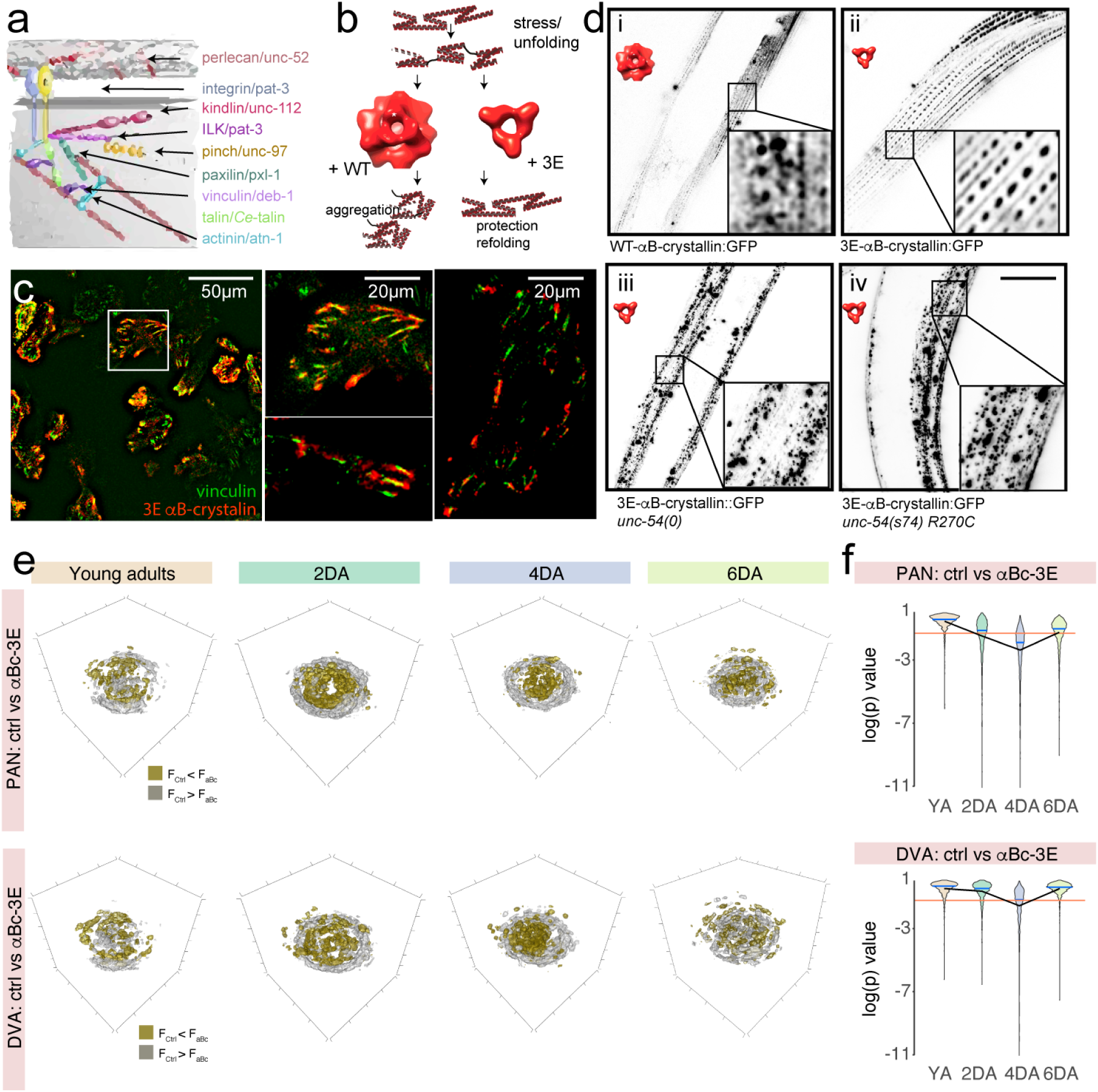
Ectopic expression of constitutive chaperones rescues proprioceptive defects. **a,** Sketch of the *C. elegans* attachment sites with similarity to focal adhesion complexes. **b,** Schematic of the *α*B-crystallin wild type and triple mutant chaperones. **c,** Representative fluorescence micrograph of BSC-1 cell culture expressing vinculin::mCherry and constitutively active 3E-*α*B-crystalin::GFP. Scale bar indicated in the figure. **d,** Representative micrograph of a transgenic animal expressing i) wild type *α*B-crystallin, ii) 3E-*α*B-crystallin, 3E-*α*B-crystallin in the body wall muscles of the iii) *unc-54(e1009)* and iv) *unc-54(s74)* mutant animals. The inset shows a magnified region of the body wall muscles in each transgenic animal. Scale bar = 40 µm. **e,** 3D plots of the statistical difference comparing the joint probability distribution (equivalent to a discrete 3D histogram) of the two forward and turning modes in the eigenworm space between N2 and age-matched tissue-specific 3E-*α*BC expression at four different ages. Silver indicates higher density for N2 control animal, golden voxels indicate higher density for tissue-specific 3E-*α*BC expression with p*<*0.01. **f,** Violin plots of the *p*-value distributions for 1000 independent tests of a bootstrapped population estimate of the 3D probability density function of an N2 control animal tested against a bootstrapped population estimate of the age matched sample from the transgenic animals expressing 3E-*α*Bc either pan-neuronally or in DVA. Orange line indicates *α*=0.05 level of significance for the hypothesis H_0_ that bootstrapped density functions derived from N2 and age animals are equal (see Methods). The blue line shows the mean p-value, the black line connects the median of the different populations.

Hence, we sought to stall or even rescue age-related changes in neuronal mechanobiology by expressing the synthetic heat-shock protein, *α*B-crystallin derived from human lens. *α*B-crystallins are ATP-independent chaperones with a strong homology to HSP-25 and whose function depend on phosphorylation. In addition, they are hypothesized to bind to mechanically stressed proteins^72, 73^ to avoid aggregation^57, 74^. Because human *α*B-crystallin resides in the cytoplasm in its inactive form, we engineered a constitutive active form of *α*B-crystallin by introducing three mutations mimicking phosphorylation (S19E, S45E, S59E; Fig. 6b). As expected, 3E*α*B crystallin localized to mechanically stressed focal adhesions in BSC-1 cells, together with the characteristic focal adhesion marker vinculin (Fig. 6c). No such colocalization is observed after treatment with latrunculinA, known to disrupt microfilament organization.

We then generated transgenic animals expressing human wild type *α*B-crystallin::GFP in body wall muscles and found a patchy localization to structures that resemble dense bodies and the endogenous HSP-25 pattern^68^ (Fig. 6d i). In line with their proposed role in chaperone activity, the constitutively phosphomimicking 3E variant localized more pronounced to the dense bodies (Fig. 6d ii). To understand if the binding of *α*B-crystallin to the dense bodies depends on force and potentially hints to mechanical unfolding, we also tested if the localization depends on muscle activity and in particular the contractile functions of muscle myosin *unc-54*. Importantly, the localization of our *α*B-crystallin reporter was drastically changed in the *unc-54(e1009)* null mutant, and was found primarily in diffuse aggregates (Fig. 6d iii), consistent with the fact that *α*B-crystallin can bind to residues that are exposed as a result of mechanical unfolding of UNC-54. This result is unlikely due to direct binding of *α*B-crystallin to UNC-54 or aberrant sarcomere assembly, as a single point mutation in UNC-54, which is not known to interfere with thick filament organization but force generation^65^, lead to the same dislocalization (Fig. 6d iv). We thus interpret these findings that 3E-*α*B-crystallin binds to mechanically strained proteins by stabilizing unfolded residues against aggregation.

The picture that is emerging suggests that mechanical stress during locomotion causes transient unfolding of the spectrin cytoskeleton, which is then cleaved by calcium-dependent proteases promoting proprioceptive decline during early age. Blocking access to exposed cleavage sites should then interfere with proteolysis and suppress this decline. We thus asked if the expression of *α*B-crystallin in DVA would lead to a detectable phenotype and suppress the age-related defects in locomotion. We tested several conditions in which we expressed the wild type and constitutively active form either pan-neuronally or specifically in DVA. Unexpectedly, in all these experimental layouts, we observed a statistically significant suppression of the age-related defect as compared to age-matched control animals (Fig. 6e,f). Intriguingly, we observed a stronger effect for the wildtype chaperone compared to the engineered 3E variant of *α*B-crystallin, both when expressed in DVA and in the whole nervous system. The underlying mechanisms remain unclear, but it due to the homology of human *α*B-crystallin to the small heat shock proteins in *C. elegans*, they may be subject to endogenous regulations and thus confer important neuroprotective role in locomotive behavior (Fig. 6e, Supplementary Fig. 7).

## 3 Discussion

Here we provided evidence that spectrin degradation leads to early onset neurodegeneration, which first becomes visible in proprioceptors and thus subtle changes in body curvature regulation. This effect can be suppressed in neuron-specific *clp-1* knockout animals but also through expression of synthetic, constitutively active heat-shock proteins. Together, our data support a model in which mechanical stresses condition the spectrin network for calcium-dependent proteolysis through calpain and probably other proteases.

Why are proprioceptors particularly vulnerable to early onset neurodegeneration? Mechanosensors in general and propriceptors in particular are subject to repetitive mechanical stresses as a part of their physiology and thus might experience non-reversible damage throughout their life. In addition, their role in detecting mechanical signals leads to reversible unfolding of load bearing molecules *AND* elevated calcium signaling as part of their force sensing function, which, in turn, may lead to unregulated activities of calcium-dependent proteases. Likewise, random oxidative modification may occur at cryptic side chains that become exposed by force have been shown to undergo accelerated aging^75^. As shown here, one of them is the ubiquitous Calpain/Calmodulin system. Thus, the coincidence of high calcium concentrations and mechanical stress may sensitize load bearing molecules of the cytoskeleton, such as the spectrin network to proteolysis at (un)folded residues. Indeed, calpain cleaves its target with a preference on buried hydrophobic residues,^55^ that may become exposed during transient unfolding, and was previously shown to act on mechanically unfolded focal adhesion kinase in cardiac muscle cells^57^. Due to their unregulated and unwanted side effects, we term this process a stochastic mechanical ‘run-on’ proteolysis, similar to how destructive run-on of wild-type biological programs causes senescent pathologies^76^. Thus, the cumulative and repetitive mechanical activity in proprioceptors increases the probability of cleavage and makes them particularly vulnerable to stress - also associated with the first phenotypes.

### Mechanical unfolding as a pathophysiological process

Several pioneering studies demonstrated that load-bearing molecules have the tendency to unfold *in vivo* when they are subjected to mechanical stresses as a part of their function^21, 24, 77, 78^. Often, these unfolded residues expose cryptic binding sites that recruit other proteins and initiate signaling cascades ^57, 58, 79–82^. Mechanical unfolding of the focal adhesion kinase at adhesion sites of cardiomyocytes was shown to recruit proteolytic enzymes that cleave FAK and lead to cardiomyocyte apoptosis. Calpain has also been shown to cleave *α*-spectrin *in vitro*, however, the significance of this cleavage site is unclear — mutant mice lacking this cleavage site in *α*-spectrin domain 10 have no discernable phenotype compared to wild type animals^83^.

Mechanical stress is a ubiquitous physical signal that continuously acts on all cells in our body - even on neurons of the central nervous system. It was shown that modest strains of 3-5% already (delivered at extremely high rates; ≈ 2 · 10^6^µm/s, ^10^) cause transient morphological alterations known as varicosities, which form with the purpose to delimit the spread of potentially cytotoxic calcium concentrations^10^. Hence, neurons appear to have evolved several mechanisms to evade the potential detrimental effect of mechanical damage, like its dissipation through controlled and reversible unfolding of tension-sensitive domains. The spectrin network, thus, acts as a shock absorber^24^ and tension homeostat, that is able to maintain a mechanical state despite being subject to massive strains in the order of 30-40% (delivered at very low rates; ≈µm/s, ^17, 23^). The transient unfolding however, with the concomitant exposure of normally buried hydrophobic residue becomes susceptible to proteolytic enzymes. This may stem from the gain in entropy associated with the release of water cage surrounding the exposed sites. Calpain, with their preference for hydrophobic residues and disordered regions may thus target them for proteolysis^53^. Intriguingly, mechanical unfolding prior cleavage may not always be detrimental but can be harnessed as a means to regulate cleavage under defined conditions^56–58^.

In future, the development of a mechanical unfolding sensor will thus be of great benefit to study the pathophysiology of compliant protein networks *in vivo* and provide mechanistic insight into the etiology of the disease.

## Acknowledgements

We would like to thank the NMSB lab members for discussion and help throughout the preparation of this work, the *Caenorhabditis* Genetic Center (CGC, supported by the National Institutes of Health, P40 OD010440) for reagents and Aleksandra Pidde for data analysis. We thank Mohamed Ali Jarboui from the Proteomic Core Facility for Medical Bioanalytics, Institute for Ophthalmic Research, Eberhard Karls University of Tübingen for performing the mass spectrometry analysis. All authors thank the Biolab and the SLN at ICFO for the use of their facility and instruments.

## Data and Code availability

All numerical data is available upon request from the corresponding author. The list of unc-70 interaction partners will be deposited in relevant databases.

## Author contribution

NM, LL, RD performed locomotion experiments, NM performed calcium imaging, RD and MK performed FRET imaging, NM, RD and AD performed the proteomic analyses, MPR, NM and RD performed molecular biology. MK and NS supervised the work and acquired funding. MK conceptualized the study. NM, MPR and MK wrote the first draft, with input from all authors.

## 5 Materials and Methods

### Animal culture and maintenance

Animals were maintained on Nematode Growth Medium (NGM) plates seeded with *Escherichia coli* OP50 bacteria. Age-synchronized young adult animals were used for all the experiments and handled as described^85^. All strains generated in this study are listed in Supplementary Data Table 3.

### Molecular biology and transgenesis

#### Insertion of mKate at the *clp-1* locus

mKate was inserted at the N-terminus using the SEC strategy^86^ to yield MSB15. To do so, two gRNAs (#6: ggtttcacagcaaaagccga; #7: aaaagccgacggaatcaaaA) were cloned into pDD162 to yield pMK106 and pMK107, respectively. These plasmids were coinjected with pMK29 harboring the SEC repair template (directing mKate to the N-terminus of the endogenous *clp-1* locus) into N2 animals and outcrossed multiple times after successful heatshock and removal of the selection cassette and expression of mKate was confirmed using confocal microscopy.

#### Insertion of loxP sites at the *clp-1* locus

After removal of the SEC cassette to generate MSB15, a single loxP site remained in the genome. This strain was used as a target to insert a second loxP site at the 3’ end of the *clp-1* endogenous locus to yield MSB973. Cloning free CRISPR tagging was employed using crRNA #67 (TATGTTTTGTTGTTTACTAT) and the ssODN HR template #42 (CATTATCTAACCAATATTTATGTTTTGTTGTTTACATAACTTCGTATAGCATACATTATACGAAGTTATTATTGGATGTGTGAAACGTATCAAATAACAGAAAA).

#### *clp-1* knockout

The *clp-1(mir60)* mutation was generated by deleting 4000 bp of the genomic coding region, using CRISPR guided by two crRNAs (GGTTTCACAGCAAAAGCCGA and TAT-GTTTTGTTGTTTACTAT). The CRISPR screening was performed using the *dpy-10* coCRISPR method^87^ into the MSB15 (mKate:clp-1) background. Jackpot broods were screened for loss of red fluorescence indicative for a successful *clp-1* deletion.

#### Tissue-specific *clp-1* knockout

The Cre-lox recombination system was employed to excise the *clp-1* gene in specific cells. Initially, the *clp-1* floxed strain (MSB973) carrying the mKate-fused endogenous *clp-1* gene was utilized. This strain was then crossed with pan-neuronal Cre (FX14125) to achieve KO of *clp-1* in the nervous system (MSB1026). For muscle-specific *clp-1* KO, a cross was performed between MSB1013 and FX16634. Similarly, for DVA-specific *clp-1* knockout, a cross was carried out between MSB1013 and MSB1030. Additional panneuronal Cre-expressing strains with *clp-1* Flox were generated using the *rgef-1* promoter, namely MSB1275, MSB1276, and MSB1277.

#### Transgenesis

Transgenesis has been performed as described before following standard procedures through microinjections and protocols described in ^88^. Random integration of extrachromosomal arrays was performed using the UV/TMP method.

#### Plasmid designs

All plasmids were design using benchling.com’s DNA editor. Synthetic or genomic DNA fragments were amplified using KOD Hot Start polymerase and assembled using Gibson’s method.

### Microscopy

#### Calcium imaging in moving animals

Calcium recordings were performed as described before^23^. In short, young adult animals were mounted onto 1-1.5% of agar pad with 3-5 µL of latex beads (Polybeads, 0.2 µm, PolySciences) to facilitate body movement and stabilize the position of field of view^22^. Calcium imaging was performed using a Leica DMi8 microscope equipped with a 25x/0.95 water immersion lens, Lumencor SpectraX LED lightsource, and a Hamamatsu Orca Flash 4 V3 sCMOS camera. The GCaMP6s calcium sensor was excited with 30% of the cyan LED of the SpectraX with a 488 nm excitation filter (≈12mW) and the calcium insensitive signal was excited with the 50% of the green/yellow LED through a 575/25 nm excitation filter (≈33mW) using a triple bandpass dichroic mirror in the filter turret (FF409/493/596-Di02-25x36, Semrock). The incident power of the excitation light was measured with a Thorlabs microscope slide power meter head (S170C) attached to PM101A power meter console. Emission was split with a Hamamatsu Gemini W-View with a 538 nm edge dichroic (Semrock, FF528-FDi1-25-36) and collected through two single band emission filters, 512/25 nm for GCaMP (Semrock, FF01-512/25-25) and 670/30 nm for mKate (Semrock, FF01-670/30-25), respectively. Both emission signals were split onto top/bottom of the image sensor, enabling differential exposure times optimized for imaging. Individual frames were acquired at 10Hz for 40-60 seconds (depending on worm movement) with an 88 ms and 50 ms exposure time, using the master pulse from the camera to trigger the light source through Hamamatsu HCImage software. Because it was impossible to resolve calcium signals throughout the long DVA axon in moving animals, we restricted our curvature analysis to the posterior region close to the cell body.

#### Image analysis

Images were processed using custom built MATLAB routines to extract the mean intensity of the cell body as a function of body centerline curvature near the tail. Due to the omnipresent autofluorescent signal in the GFP channel, adaptive thresholding was able to separate the animal backbone from the background signal. First, the raw images from the GCaMP channel were binarized and eroded to find the skeleton of the worm in each frame. An iterative approach was chosen to prune the branches to get the longest path describing the centerline of the worm^89^. Next, a segment of the centerline enough to capture one bend of the worm in the tail region was chosen, which was further divided into two equal segments using three points. These three points were used to construct a triangle, and subsequently a circumcircle. Finally, the curvature is calculated at the middle point of the three points using the radius of the circumcircle by the formula *κ* = 1/R, where R is the radius of the circumcircle and *κ* is the curvature, and directionality of the curvature was determined by the sign of the tangential angle at the point where the curvature was calculated.

The neuron was labelled manually for the first frame in the mKate, calcium insensitive channel, and automatically tracked in subsequent frames, based on a local search in the vicinity of the location in the previous frames. The area, position and intensity was collected in the mKate channel and its position was mapped onto the GCaMP channel to extract the intensity in each frame. Finally, the calcium sensitive signal was divided by the insensitive signal after background correction (for Fig. S4A,C) to obtain the ratio *R* which was normalized to the baseline ratio *R*_0_. The trough of the periodic signals was taken as *R*_0_. The background subtracted calcium traces were divided by the background subtracted mKate signal to yield the ratiometric intensity signal:

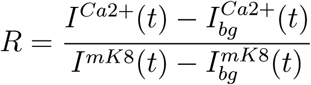

#### Confocal imaging

Fluorescence images were taken using an inverted confocal microscope (Nikon Ti2 Eclipse) with a 60x/1.4 NA oil immersion lens. Animals were imaged live in 3 mM levamisole on 5-6% agarose pads. mCherry was excited using the 561 nm laser, 20-30% power intensity and transmitted through a 594 nm emission filter. Exposure time was 100-200 ms, depending on the strain to image. GFP was excited with a 488 nm laser, 20-40% power intensity and transmitted through a 521 nm emission filter. Exposure time was 100-200 ms.

For CLP-1 expression, MSB15 transgenic worms were synchronized and cultured on NGM/OP50 medium before the imaging procedure. Samples were prepared by placing the worms on a 2% agar pad containing 3mM levamisole. Snapshots using the fluorescent channel in the head, mid-body, and tail regions were taken. The fluorescent signal was then compared between groups ranging from the young adult stage to the day-12 adult stage.

#### FRET imaging

FRET imaging was performed on a Leica DMI6000 SP5 confocal microscope using the 63x/1.4 NA oil immersion lens using the sensitized emission approach. As described in detail in ^35^, three images were collected: the direct donor (mTFP2) excitation and emission, donor excitation and acceptor emission and direct acceptor (mVenus) excitation and emission. mTFP2 was excited with 458 nm (≈9 µW), mVenus with 514 nm (≈4 µW) line of an Argon ion laser at 80% and 11% transmission respectively (25% power). The incident power of the excitation light was measured with a Thorlabs microscope slide power meter head (S170C) attached to PM101A power meter console. The FRET index was extracted as described^35^. The piece-wise linear fit in Fig. 5h was performed using the `segmented` package in R. The breakpoint was determined at ≈6 days.

#### Locomotion analysis

Recordings of locomotion behavior and analysis was performed exactly as described in detail in Reference ^23^. Briefly, 3-5 min long videos of young adult synchronized animals transferred to non-seeded NGM plates were recorded at 25 fps using a home-built tracking platform. Imaging processing was done by using eigenworms with custom MATLAB scripts. To compare the genotypes, animal locomotion behavior was decomposed using the Eigenworm analysis^26^. To construct and visualize the 3D distributions of the modes (*a*_1_, *a*_2_, *a*_3_), the 3D kernel density estimate of the first three modes was calculated in R using the `ks` package^90^ with an unconstrained plug-in selector bandwidth. We choose to indicate the 10, 25 and 50% contours of the highest density regions in the manifold and the 2D projections of the floating data cloud along the corresponding planes. To compare two different data sets and test for the null hypothesis that the two kernel density functions are similar, we resampled the highly oversampled population by bootstrapping to avoid spurious significance due to long tailed outliers. The resampling and testing was performed 1000 times to yield a distribution of *p*-values which is displayed as a violin plot summarizing each figure of locomotion data. Importantly, the resampling does not lead to significant discrimination of the downsampled and original dataset within the same population.

For the final quantification and display of the data, two 3D distributions of the Eigenworms in the kde space were compared. Only voxels in which a statistically significant difference was calculated were displayed in Fig. 1a i; 5a, e and Fig. 6. Exact number of animals and frames collected for the analysis can be found in Supplementary Table 2

### Western Blot of UNC-70

#### Worm lysis procedure

Approximately 5,000-10,000 age-synchronized adult worms were grown onto peptone-enriched plates supplied by NA22 *E. coli*. Worms were rinsed repeatedly with cold distilled water until the solution was devoid of bacterial cells. In the final wash, excess water was removed, and the pellet was then transferred to a 15-mL tube and stored in -80°C freezer. To initiate lysis, the frozen pellet was thawed and mixed with an equal volume of the 2x lysis buffer (50mM Tris, 300mM NaCl, 2% Triton X-100, phosphate and protease inhibitors (Sigma references P5726 and P8340 respectively)). Subsequently, a clean magnetic stir bar was placed in the tube, which was then subjected to vibrations on a platform (vortex) until the pellet was completely disrupted, resulting in a clear solution. This process was carried out while maintaining the temperature on ice. The lysate was carefully transferred to a 2-mL tube kept on ice. The obtained lysate was subjected to three rounds of sonication, each lasting 30 seconds, and subsequently centrifuged at 14.000 rpms on a tabletop centrifuge (4°C) for 15 minutes. The supernatant, now clear of debris, was collected into a fresh tube and stored at -20°C for future use.

The total protein concentration in the sample was assessed using the Bradford assay with Bovine serum albumin (BSA) as the protein standard.

#### Polyacrylamide Gel Electrophoresis (SDS-PAGE)

The protein lysate was mixed with a 5x loading buffer consisting of 250 mM Tris (pH 6.8), 12.5 mM EDTA, 10% SDS, 25% glycerol, 8M urea, 20 mM DTT, and 0.025% Bromphenol blue. Gel electrophoresis was conducted using a Biorad gel running apparatus. The samples were loaded into the wells of a 4–15 % gradient gel (Biorad, #4561085DC) and run at 300V and 75mA. Subsequently, the gel was carefully removed from the mold, and transferred to 0.45 *µ*m nitrocellulose Membrane, (Biorad, #1620145), at 100V for 2 hours within an ice block. The whole transferred protein was quantified with Ponceau to verify transfer and quantify whole protein amount.

#### Primary antibody incubation

After Ponceau staining, the membrane was washed with TBST buffer to remove any residual dye. To block non-specific protein binding, the nitrocellulose membrane was soaked in TBST with 5% skim milk at room temperature for 1 hour. Subsequently, the primary antibody solution (Anti-GFP antibody, Abcam, ab13970) was applied to the membrane at room temperature for 1 hour on a rocking platform. The membrane was then quickly rinsed with 20 mL of TBST buffer, followed by three washes with TBST on the rocking platform for 10 minutes each.

#### Secondary antibody incubation

The primary antibody on the membrane was subsequently probed with a secondary antibody against chicken protein (IRDye® 800CW Donkey anti-Chicken Secondary Antibody, Licor). The secondary antibody solution was prepared by mixing the antibody with TBST + 5% skim milk (diluted 1:15000) and applied to the membrane. The incubation was carried out at room temperature on a rocking platform for 1 hour. After the incubation, the antibody solution was discarded, and the membrane was quickly rinsed with TBST buffer. Subsequently, the membrane was washed three times with TBST buffer for 10 minutes each on the rocking platform.

#### Imaging

The western blot membrane was imaged using the Azure c600 device. The fluorescent imaging mode was selected, and the auto-exposure time was used to capture the image. The protein marker was captured using visible light. The quantification of the protein bands on the western blot was performed by comparing the intensity of the bands.

### UNC-70::GFP CoIP and Mass spectrometry analysis

#### Worm lysis procedure

Frozen worm pellets were thawed on ice, mixed with an equal volume of the 2x lysis buffer (50 mM Tris, 300 mM NaCl, and 2 % Triton X-100) and incubated for 30 min on ice. The lysate was then mechanically homogenized using a T10 basic Ultra-Turrax (IKA) and centrifuged at high speed (19,000 x*g*) at 4 °C for 30 minutes. The supernatant, which contains cytoplasmic and membrane proteins, was transferred into a fresh tube and immediately used for co-immunoprecipitation.

Co-immunoprecipitation was performed using GFP-Trap Magnetic Agarose beads (Chromotek, gtma-100). According to manufacturer’s protocol, the beads were equilibrated by washing three times with 500 µL ice-cold dilution buffer. 1 mL of lysate was then added to 50 µL washed beads, and incubated at 4 °C for 1 hour while rotating. The supernatant containing the non-bound fraction was removed and beads were washed three times with wash buffer. One set of beads was re-suspended in dilution buffer and used for mass spectrometry analysis. Another set of beads was re-suspended in 100 µL sample buffer and boiled for 5 min at 95 °C to dissociate immunocomplexes from the beads. The supernatant was analyzed by western blot as described above using chicken@GFP (Abcam, ab13970, 1:2000) and rabbit@mKate (Origene, TA150072; 1:1000).

#### Mass spectrometry analysis

Mass spectrometry has been performed as previously described^91^. In brief, Mass spectrometry (MS) analysis was conducted using an Ultimate3000 RSLC system connected to an Orbitrap Fusion Tribrid mass spectrometer (Thermo Fisher Scientific). Tryptic peptides were first loaded onto a µPAC Trapping Column (pillar diameter 5 µm, an inter-pillar distance of 2.5 µm, a pillar length/bed depth of 18 µm, an external porosity of 9%, a bed channel width of 2 mm, and a length of 10 mm). The loading flow rate was 10 µl per minute in 0.1% trifluoroacetic acid in HPLC-grade water.

Peptides were then eluted and separated on a PharmaFluidics µPAC nano-LC column (50 cm µPAC C18), which had a pillar diameter of 5 µm, an inter-pillar distance of 2.5 µm, a pillar length/bed depth of 18 µm, an external porosity of 59%, a bed channel width of 315 µm, and a bed length of 50 cm. The pillars were superficially porous with a shell thickness of 300 nm and pore sizes between 100 and 200 Å. Separation was performed using a linear gradient from 2% to 30% buffer B (80% acetonitrile, 0.08% formic acid in HPLC-grade water) in buffer A (2% acetonitrile, 0.1% formic acid in HPLC-grade water) at a flow rate of 300 nl per minute. The remaining peptides were eluted using a short gradient from 30% to 95% buffer B, with a total gradient run time of 120 minutes.

The MS parameters were as follows: full MS spectra were acquired within a scan range of 335–1,500 m/z with a resolution of 120,000 at m/z 200. MS/MS data were acquired in top speed mode with a 3-second cycle time. The maximum injection time was 50 ms, with an AGC target of 400,000 and an isolation window of 1.6 m/z. Ions with charge states between 2 and 7 were fragmented sequentially using higher-energy collisional dissociation.

Analysis was performed using the MaxQuant software as described^91^.

## 6 Supplementary Figures

**Supplementary Fig. 1.**
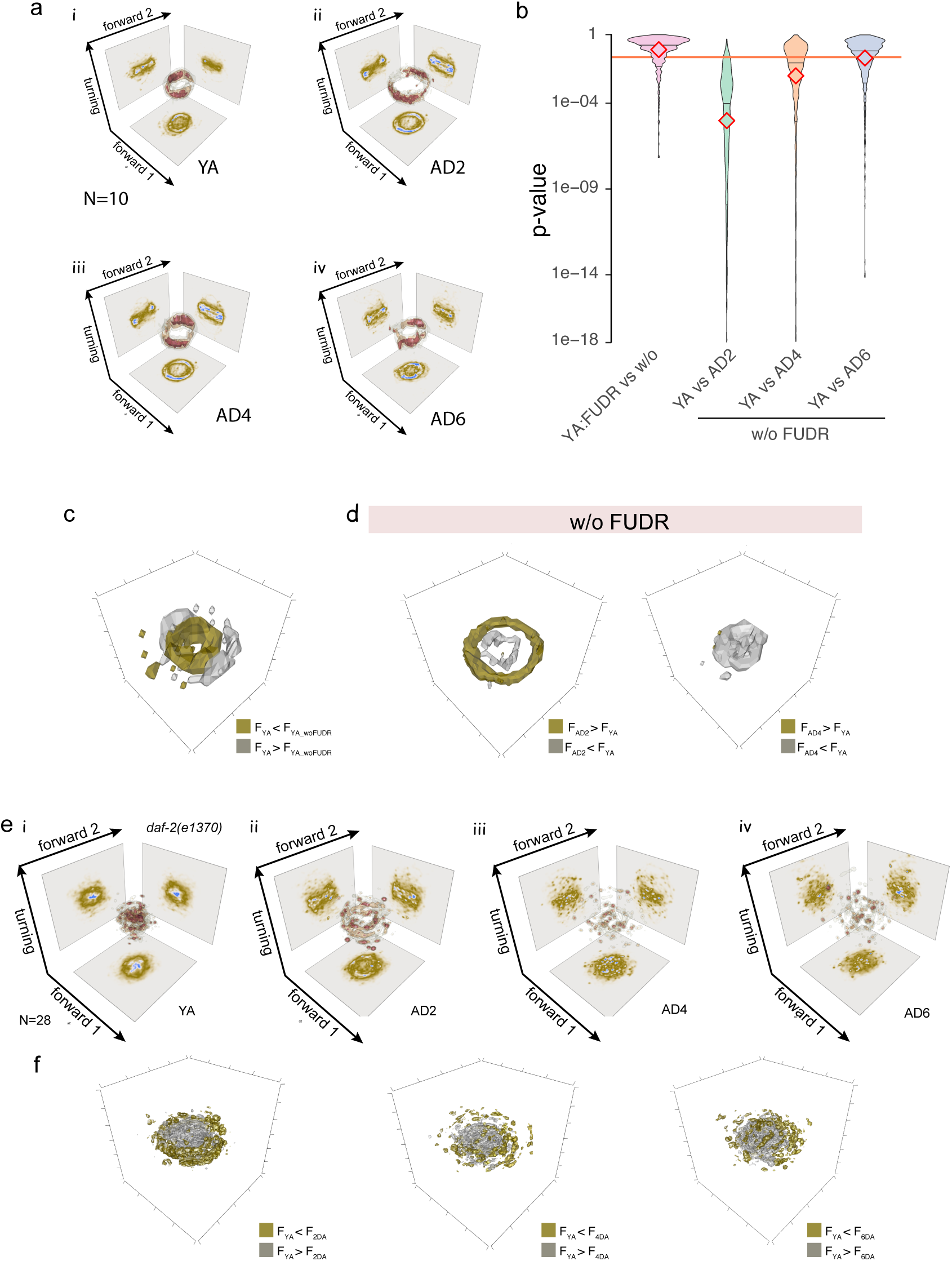
Changes in animal locomotion are independent from FUDR. **a** 3D kernel density estimate for the joint probability distribution (equivalent to a discrete 3D histogram) of the two forward and turning modes in the eigenworm space for i) young adults, ii) adult day 2, iii) adult day 4 and iv) adult day 6. Color scale = (brown, low density; red, high density). **b** Violin plots of the *p*-value distributions for 1000 independent tests of a bootstrapped population estimate of control 3D probability density function (N2 YA) tested against a bootstrapped population estimate of the aged sample. Orange line indicates *α*=0.05 level of significance for the hypothesis H_0_ that bootstrapped density functions derived from N2 and age animals are equal (see Methods). The red line shows the mean p-value, diamonds = median. **c** Statistical test comparing the local kernel density function *F* of young adult animals with and without FUDR. YA animals with FUDR lead to a slightly larger body bends, indicated by the silver voxels surrounding the golden ones. **d** Statistical test comparing the local kernel density function *F* of young adults and older (AD2 and AD4) animals. The test is performed on each voxel in a 100x100x100 grid (see Methods) and color coded according to its outcome. Golden voxels indicates that AD2 is significantly larger (gold) or smaller (silver) than young adults. Voxels with p*>*0.01 are not shown. **e** 3D kernel density estimate for the joint probability distribution (equivalent to a discrete 3D histogram) of the two forward and turning modes in the eigenworm space for i) young adults, ii) adult day 2, iii) adult day 4 and iv) adult day 6 of daf-2 mutant animals. Color scale = (brown, low density; red, high density). **f** Statistical test comparing the local kernel density function *F* of young adults and older (AD2, AD4 and AD6) animals. The test is performed on each voxel in a 100x100x100 grid (see Methods) and color coded according to its outcome. Golden voxels indicates that older animals in the daf-2 background crawl with significantly larger curvatures than young adults. This observation is similar to control animals. Voxels with p*>*0.01 are not shown.

**Supplementary Fig. 2.**
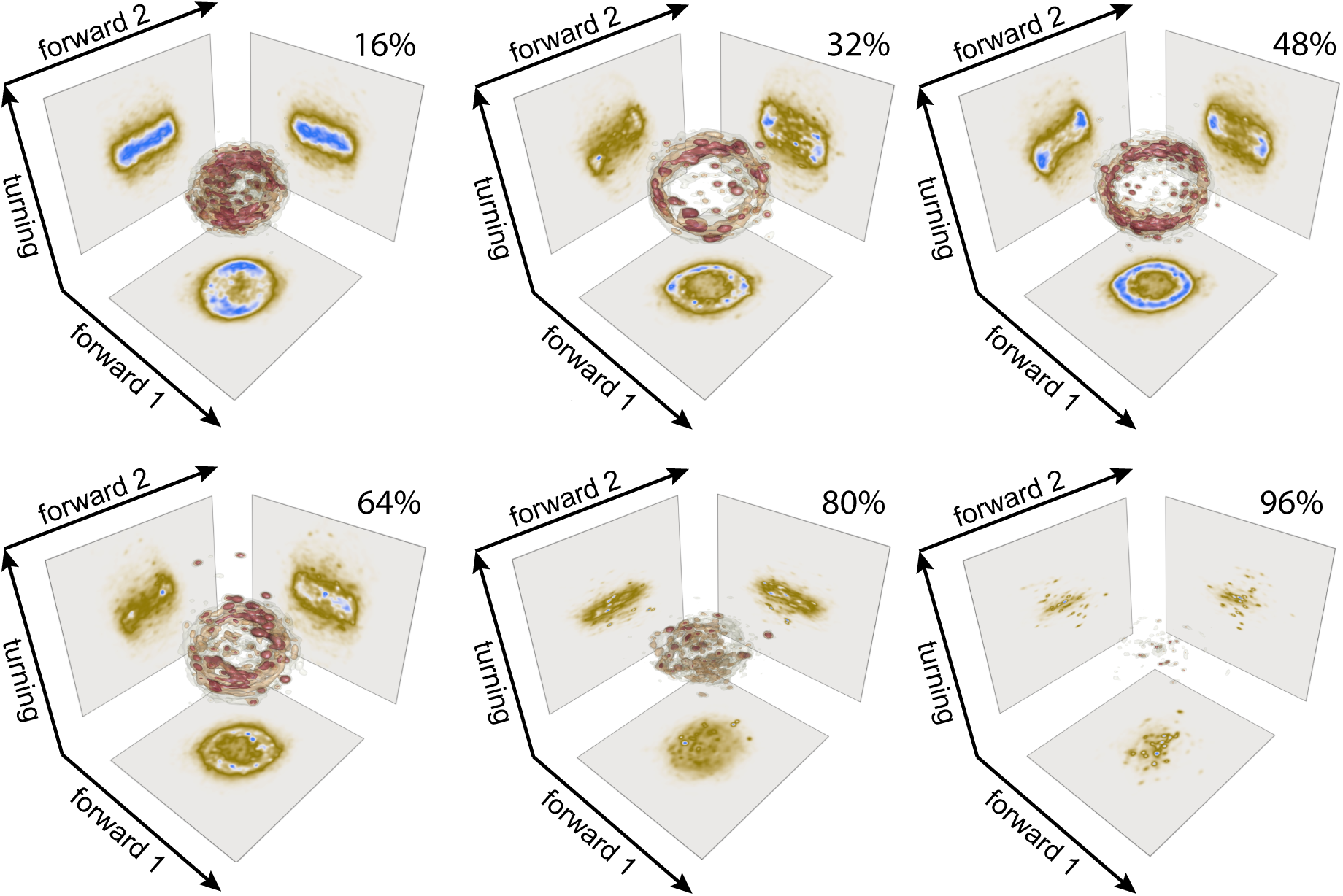
Eigenworms according to the relative age. 3D kernel density estimate for the joint probability distribution (equivalent to a discrete 3D histogram) of the two forward and turning modes in the eigenworm space for 6 different categories spanning the indicated % of their total lifespan. Color scale = (brown, low density; red, high density).

**Supplementary Fig. 3.**
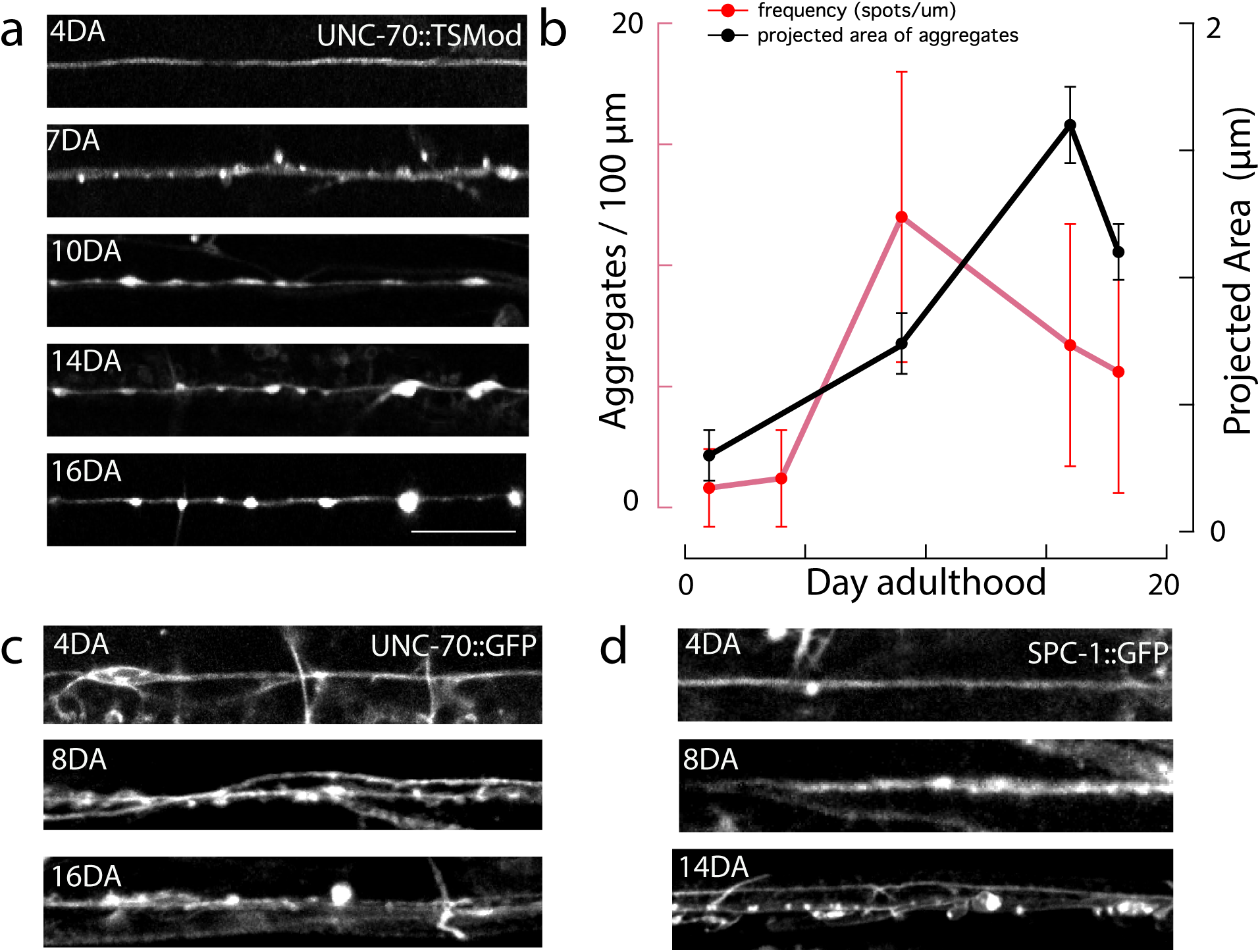
Animal age leads to aggregates in the spectrin network. **a,** Representative image of progressively aged animals from adult day 4 through adult day 16. **b,** Quantification of the aggregate size (black line) and the spacing between the aggregates (red line) as a function of animal age. Mean±SD. **c,** Representative images of different neurons expressing a single copy knock-in of GFP at the endogenous *unc-70* locus for three different ages. **d,** Representative images of different neurons expressing a single copy knock-in of GFP at the endogenous *spc-1* locus for three different ages. Scale bar for all images = 10 µm.

**Supplementary Fig. 4.**
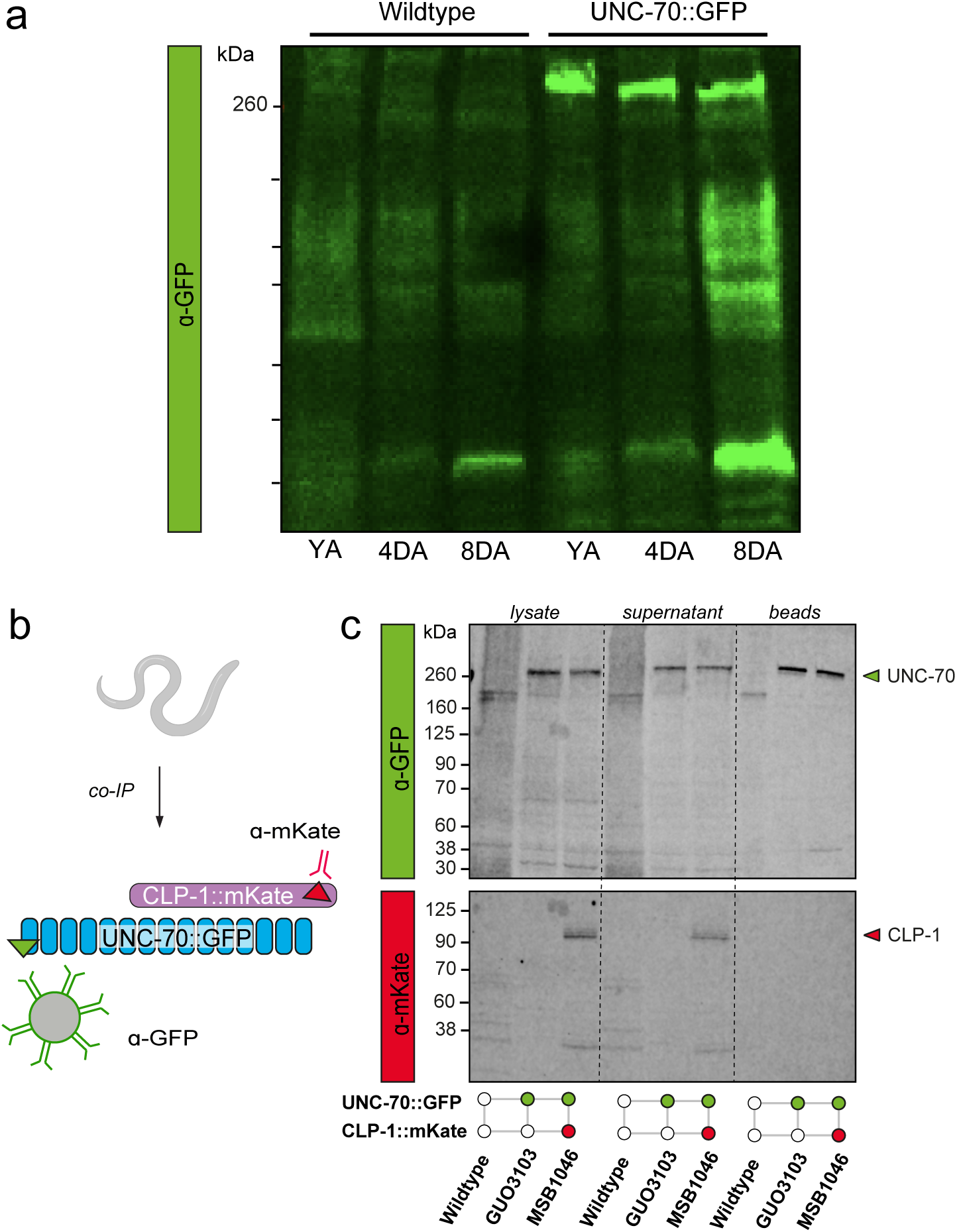
Co-Immunoprecipitation of CLP-1::mKate with UNC-70::GFP. **a,** Western blot of unlabeled wild type controls and UNC-70:GFP transgenic worms show no unspecific GFP-labeling in the untagged samples. **b,** Schematic of co-IP of UNC-70 and CLP-1 transgenic worms. **c,** Immunoblot of co-IP using UNC-70 as bait to capture CLP-1. UNC-70 (green arrowhead) and CLP-1 (red arrowhead) were detected using anti-GFP and anti-mKate antibodies, respectively. Blot shows lysate of worms before co-IP (’lysate’), supernatant from beads after co-IP (’supernatant’) and the bound fraction from beads (’beads’).

**Supplementary Fig. 5.**
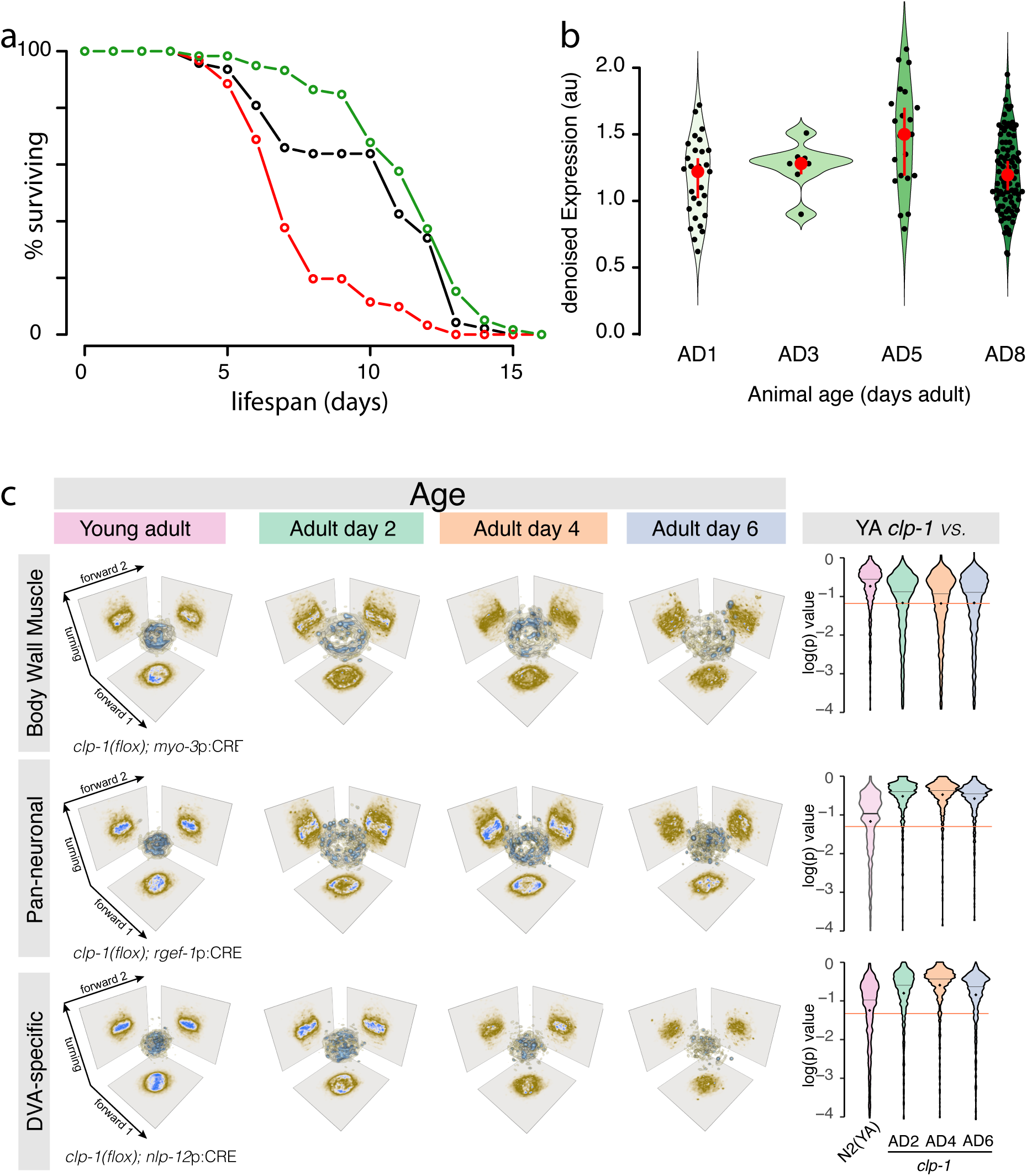
Tissue-specific calpain deletion results in defects in animal lifespan. **a,** Lifespan curves for control N2 animals (black), *clp-1* knockout (red) and *clp-1* overexpression animals (green). **b,** Expression of *clp-1* in DVA in log2 (count per 10k) at each time point represented as a violin plot ^63^. Expression increases until day 6 and then declines **c,d,** Panneuronal and DVA knockout suppress the locomotion defect during early age. (c) 3D plot of the estimate for the joint probability distribution (equivalent to a discrete 3D histogram) of the two forward and turning modes in the eigenworm space for the tissue-specific *clp-1* mutations in i) all neurons, ii) the body wall muscles and iii) specifically in DVA at four different ages. Color scale = (brown, low density; blue, high density). (d) Violin plots of the *p*-value distributions for 1000 independent tests of a bootstrapped population estimate of control 3D probability density function (N2 YA) tested against a bootstrapped population estimate of the aged sample. Orange line indicates *α*=0.05 level of significance for the hypothesis H_0_ that bootstrapped density functions derived from N2 and old animals are equal (see Methods). The red line shows the mean p-value, diamonds = median.

**Supplementary Fig. 6.**
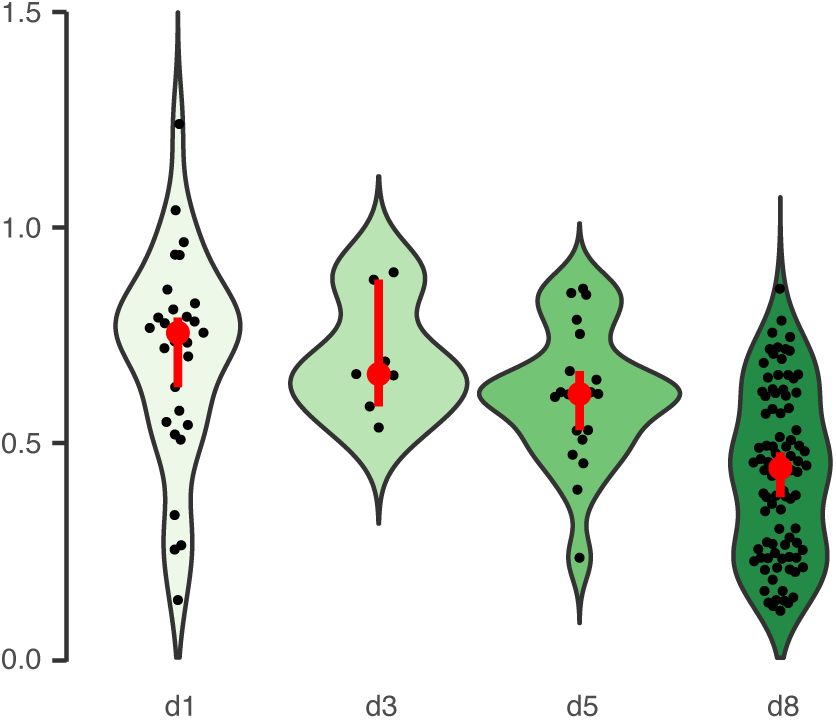
Expression of *hsp-25* vs age. Expression of *hsp-25* in DVA in log2 (count per 10k) at each time point represented as a violin plot.

**Supplementary Fig. 7.**
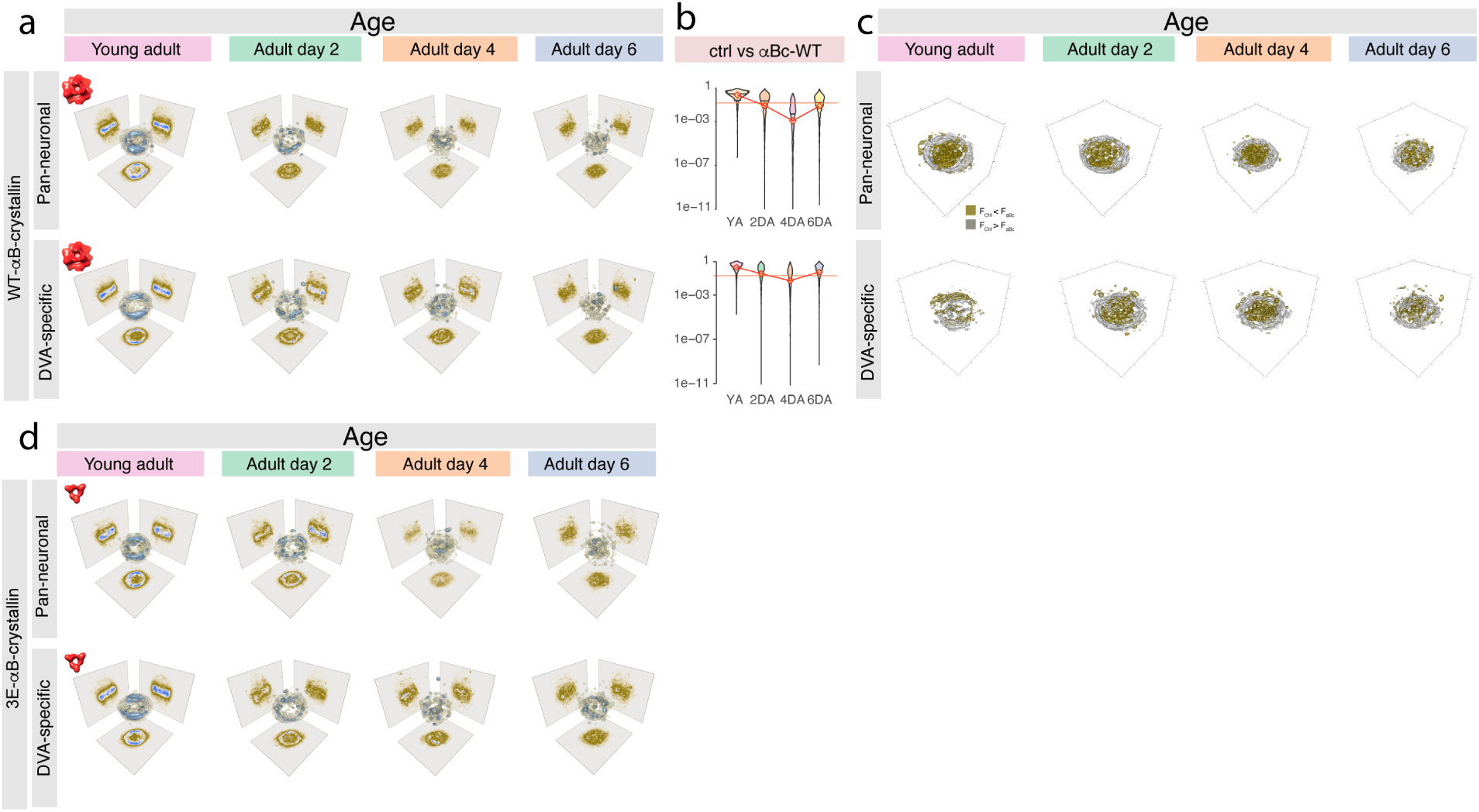
Expression of *α*B-crystallin in different tissues differently suppress locomotion defect during early age. **a-c** Quantification of the locomotion behavior for animals expressing the wild type *α*B-Crystallin. **a,** 3D plot of the estimate for the joint probability distribution (equivalent to a discrete 3D histogram) of the two forward and turning modes in the eigenworm space for animals expressing wild type *α*B-Crystallin pan-neuronally or in DVA exclusively for 4 different time points. Color scale = (brown, low density; blue, high density). **b,** Violin plots of the *p*-value distributions for 1000 independent tests of a bootstrapped population estimate of the 3D probability density function of an N2 control animal tested against a bootstrapped population estimate of the age matched sample from the transgenic animals expressing wt-aBC either pan-neuronally or in DVA. Orange line indicates *α*=0.05 level of significance for the hypothesis H_0_ that bootstrapped density functions derived from N2 and age animals are equal (see Methods). The black line shows the mean p-value, diamonds indicate the median. **c,** 3D plot of the statistical difference comparing the joint probability distribution (equivalent to a discrete 3D histogram) of the two forward and turning modes in the Eigenworm space between N2 and age-matched tissuespecific 3E-aBC expression at four different ages. Silver indicates higher density for N2 control animal, golden voxels indicate higher density for tissue-specific 3E-aBC expression with p*<*0.01. **d,** 3D plot of the estimate for the joint probability distribution (equivalent to a discrete 3D histogram) of the two forward and turning modes in the eigenworm space for animals expressing constitutively active 3E *α*B-crystallin i) pan-neuronally or ii) in DVA exclusively for 4 different time points. Color scale = (brown, low density; blue, high density).

## 7 Supplementary Videos

**Supplementary Video 1** Representative video of a one day old adult animal expressing GCaMP6s in DVA, moving its tail under the coverslip. Scale bar = 40µm.

**Supplementary Video 2** Representative video of a six day old adult animal expressing GCaMP6s in DVA, moving its tail under the coverslip. Scale bar = 40µm.

## 8 Supplementary Tables

**Table 1.** Mass spectrometry results from the UNC-70::GFP pulldown assay.

**Table 2.** Number of data points for different genotypes used analyzing the animal locomotion behavior.

| Strain | genotype | age | Fig. | drug? | worms | frames |
| --- | --- | --- | --- | --- | --- | --- |
| N2 | ctrl | YA | 1 | FUDR | 45 | 196158 |
| N2 | ctrl | AD2 | 1 | FUDR | 45 | 192306 |
| N2 | ctrl | AD4 | 1 | FUDR | 45 | 169209 |
| N2 | ctrl | AD6 | 1 | FUDR | 45 | 167782 |
| N2 | ctrl | AD8 | 1 | FUDR | 45 | 167788 |
| N2 | ctrl | AD10 | 1 | FUDR | 41 | 93072 |
| N2 | ctrl | AD12 |  | FUDR | 20 | 37250 |
| N2 | ctrl | AD14 | 1 | FUDR | 9 | 22341 |
| N2 | ctrl | AD16 | 1 | FUDR | 5 | 8867 |
| N2 | ctrl | YA | S1 | w/o FUDR | 10 | 15000 |
| N2 | ctrl | AD2 | S1 | w/o FUDR | 10 | 15000 |
| N2 | ctrl | AD4 | S1 | w/o FUDR | 10 | 15000 |
| N2 | ctrl | AD6 | S1 | w/o FUDR | 10 | 15000 |
| CB1370 | *daf-2* | YA | S1 | FUDR | 28 | 117629 |
| CB1370 | *daf-2* | AD2 | S1 | FUDR | 28 | 97974 |
| CB1370 | *daf-2* | AD4 | S1 | FUDR | 28 | 99148 |
| CB1370 | *daf-2* | AD6 | S1 | FUDR | 28 | 116036 |
| MSB1031 | *clp-1(mir60)* | YA | 5b | FUDR | 20 | 81538 |
| MSB1026 | *clp-1(mir1mir58), tmls777* | YA | 5e | FUDR | 78 | 322405 |
| MSB1026 | *clp-1(mir1mir58), tmls777* | AD2 | 5e | FUDR | 78 | 316336 |
| MSB1026 | *clp-1(mir1mir58), tmls777* | AD4 | 5e | FUDR | 77 | 316336 |
| MSB1026 | *clp-1(mir1mir58), tmls777* | AD6 | 5e | FUDR | 72 | 262844 |
| MSB1039 | *clp-1(mir1mir58), tmls1087* | YA | 5e | FUDR | 20 | 71173 |
| MSB1039 | *clp-1(mir1mir58), tmls1087* | AD2 | 5e | FUDR | 20 | 86014 |
| MSB1039 | *clp-1(mir1mir58), tmls1087* | AD4 | 5e | FUDR | 20 | 83509 |
| MSB1039 | *clp-1(mir1mir58), tmls1087* | AD6 | 5e | FUDR | 20 | 82114 |
| MSB1040 | *clp-1(mir1mir58), mirls71* | YA | 5e | FUDR | 40 | 172427 |
| MSB1040 | *clp-1(mir1mir58), mirls71* | AD2 | 5e | FUDR | 40 | 130727 |
| MSB1040 | *clp-1(mir1mir58), mirls71* | AD4 | 5e | FUDR | 40 | 136179 |
| MSB1040 | *clp-1(mir1mir58), mirls71* | AD6 | 5e | FUDR | 38 | 124236 |
| MSB1049 | *mirEx440[rab-3p::abC-WT]* | YA | 6e; S6a | FUDR | 20 | 87912 |
| MSB1049 | *mirEx440[rab-3p::abC-WT]* | AD2 | 6e; S6a | FUDR | 20 | 73683 |
| MSB1049 | *mirEx440[rab-3p::abC-WT]* | AD4 | 6e; S6a | FUDR | 20 | 87547 |
| MSB1049 | *mirEx440[rab-3p::abC-WT]* | AD6 | 6e; S6a | FUDR | 20 | 83493 |
| MSB1053 | *mirEx444[nlp-12p::abC-WT]* | YA | 6e; S6a | FUDR | 20 | 80212 |
| MSB1053 | *mirEx444[nlp-12p::abC-WT]* | AD2 | 6e; S6a | FUDR | 20 | 86847 |
| MSB1053 | *mirEx444[nlp-12p::abC-WT]* | AD4 | 6e; S6a | FUDR | 20 | 85480 |
| MSB1053 | *mirEx444[nlp-12p::abC-WT]* | AD6 | 6e; S6a | FUDR | 20 | 84016 |
| MSB1055 | *mirEx446[rab-3p::abC-3E]* | YA | 6e; S6a | FUDR | 19 | 68014 |
| MSB1055 | *mirEx446[rab-3p::abC-3E]* | AD2 | 6e; S6a | FUDR | 19 | 80900 |
| MSB1055 | *mirEx446[rab-3p::abC-3E]* | AD4 | 6e; S6a | FUDR | 19 | 99148 |
| MSB1055 | *mirEx446[rab-3p::abC-3E]* | AD6 | 6e; S6a | FUDR | 19 | 74026 |
| MSB1060 | *mirEX451[nlp-12p::abC-3E]* | YA | 6e; S6a | FUDR | 20 | 82965 |
| MSB1060 | *mirEX451[nlp-12p::abC-3E]* | AD2 | 6e; S6a | FUDR | 20 | 87425 |
| MSB1060 | *mirEX451[nlp-12p::abC-3E]* | AD4 | 6e; S6a | FUDR | 20 | 86752 |
| MSB1060 | *mirEX451[nlp-12p::abC-3E]* | AD6 | 6e; S6a | FUDR | 20 | 83936 |

**Table 3.**
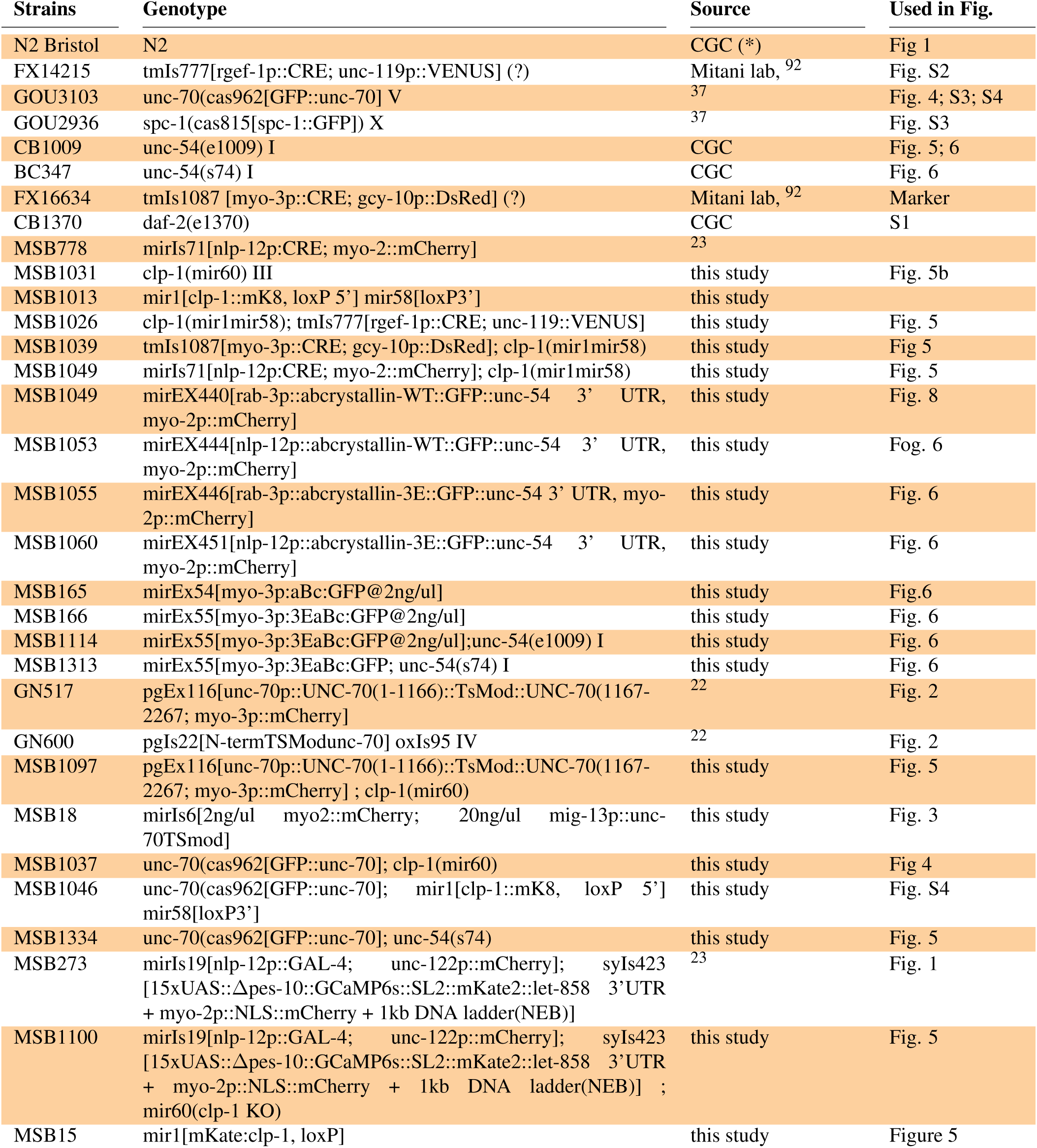
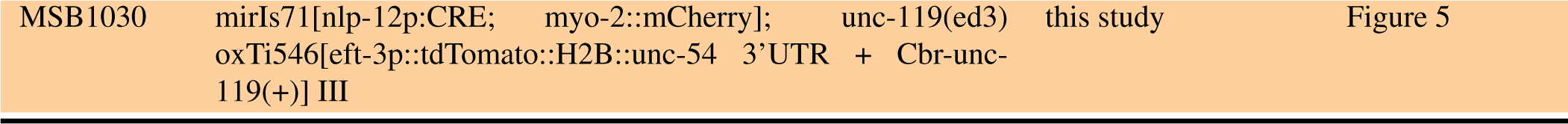
Strains used in this study.

| Strains | Genotype | Source | Used in Fig. |
| --- | --- | --- | --- |
| N2 Bristol | N2 | CGC (*) | Fig 1 |
| FX14215 | tmIs777[rgef-1p::CRE; unc-119p::VENUS] (?) | Mitani lab,<sup>92</sup> | Fig. S2 |
| GOU3103 | unc-70(cas962[GFP::unc-70] V | <sup>37</sup> | Fig. 4; S3; S4 |
| GOU2936 | spc-1(cas815[spc-1::GFP]) X | <sup>37</sup> | Fig. S3 |
| CB1009 | unc-54(e1009) I | CGC | Fig. 5; 6 |
| BC347 | unc-54(s74) I | CGC | Fig. 6 |
| FX16634 | tmIs1087 [myo-3p::CRE; gcy-10p::DsRed] (?) | Mitani lab,<sup>92</sup> | Marker |
| CB1370 | daf-2(e1370) | CGC | S1 |
| MSB778 | mirIs71[nlp-12p:CRE; myo-2::mCherry] | <sup>23</sup> |  |
| MSB1031 | clp-1(mir60) III | this study | Fig. 5b |
| MSB1013 | mir1[clp-1::mK8, loxP 5'] mir58[loxP3'] | this study |  |
| MSB1026 | clp-1(mir1mir58); tmIs777[rgef-1p::CRE; unc-119::VENUS] | this study | Fig. 5 |
| MSB1039 | tmIs1087[myo-3p::CRE; gcy-10p::DsRed]; clp-1(mir1mir58) | this study | Fig 5 |
| MSB1049 | mirIs71[nlp-12p:CRE; myo-2::mCherry]; clp-1(mir1mir58) | this study | Fig. 5 |
| MSB1049 | mirEX440[rab-3p::abcrystallin-WT::GFP::unc-54 3' UTR, myo-2p::mCherry] | this study | Fig. 8 |
| MSB1053 | mirEX444[nlp-12p::abcrystallin-WT::GFP::unc-54 3' UTR, myo-2p::mCherry] | this study | Fog. 6 |
| MSB1055 | mirEX446[rab-3p::abcrystallin-3E::GFP::unc-54 3' UTR, myo-2p::mCherry] | this study | Fig. 6 |
| MSB1060 | mirEX451[nlp-12p::abcrystallin-3E::GFP::unc-54 3' UTR, myo-2p::mCherry] | this study | Fig. 6 |
| MSB165 | mirEx54[myo-3p:aBc:GFP@2ng/ul] | this study | Fig.6 |
| MSB166 | mirEx55[myo-3p:3EaBc:GFP@2ng/ul] | this study | Fig. 6 |
| MSB1114 | mirEx55[myo-3p:3EaBc:GFP@2ng/ul];unc-54(e1009) I | this study | Fig. 6 |
| MSB1313 | mirEx55[myo-3p:3EaBc:GFP; unc-54(s74) I | this study | Fig. 6 |
| GN517 | pgEx116[unc-70p::UNC-70(1-1166)::TsMod::UNC-70(1167-2267; myo-3p::mCherry] | <sup>22</sup> | Fig. 2 |
| GN600 | pgIs22[N-termTSMoDunc-70] oxIs95 IV | <sup>22</sup> | Fig. 2 |
| MSB1097 | pgEx116[unc-70p::UNC-70(1-1166)::TsMod::UNC-70(1167-2267; myo-3p::mCherry] ; clp-1(mir60) | this study | Fig. 5 |
| MSB18 | mirIs6[2ng/ul myo2::mCherry; 20ng/ul mig-13p::unc-70TSMoD] | this study | Fig. 3 |
| MSB1037 | unc-70(cas962[GFP::unc-70]; clp-1(mir60) | this study | Fig 4 |
| MSB1046 | unc-70(cas962[GFP::unc-70]; mir1[clp-1::mK8, loxP 5'] mir58[loxP3'] | this study | Fig. S4 |
| MSB1334 | unc-70(cas962[GFP::unc-70]; unc-54(s74) | this study | Fig. 5 |
| MSB273 | mirIs19[nlp-12p::GAL-4; unc-122p::mCherry]; syIs423 [15xUAS::Δpes-10::GCaMP6s::SL2::mKate2::let-858 3'UTR + myo-2p::NLS::mCherry + 1kb DNA ladder(NEB)] | <sup>23</sup> | Fig. 1 |
| MSB1100 | mirIs19[nlp-12p::GAL-4; unc-122p::mCherry]; syIs423 [15xUAS::Δpes-10::GCaMP6s::SL2::mKate2::let-858 3'UTR + myo-2p::NLS::mCherry + 1kb DNA ladder(NEB)] ; mir60(clp-1 KO) | this study | Fig. 5 |
| MSB15 | mir1[mKate:clp-1, loxP] | this study | Figure 5 |
| MSB1030 | mirIs71[nlp-12p:CRE; myo-2::mCherry]; unc-119(ed3)<br>oxTi546[eft-3p::tdTomato::H2B::unc-54 3'UTR + Cbr-unc-119(+)] III | this study | Figure 5 |

